# Engineering the *Lr67*-associated G144R substitution by prime editing enhances leaf rust resistance in wheat

**DOI:** 10.64898/2026.11.12.765645

**Authors:** Joshua Waites, Rafael E. Venado, Easter D. Syombua, Ana Isabel Rita Castillo, Jose Pablo Espejel, Laurajean Lewis, Flavio Breseghello, Sarah Hearne, Anindya Bandyopadhyay

## Abstract

Wheat rust diseases threaten global wheat production, highlighting the need for durable sources of resistance. The Lr67-associated G144R substitution in the sugar transport protein gene TaSTP13-D1 confers broad-spectrum resistance to multiple fungal pathogens. Here, we used an enhanced prime-editing system (ePPEplus) to introduce this resistance-associated substitution into the three TaSTP13 homoeologs in hexaploid wheat. Using the wheat-compatible JWPE1 vector, we achieved editing frequencies of 84 to 92% across the A, B, and D genomes in regenerated T0 plants. Homozygous edits were recovered in all three homoeologs, including one line carrying homozygous edits across all three genomes. Disease assays showed that edited lines had reduced leaf rust severity and fungal biomass compared with susceptible controls. Notably, the line carrying edits in all three homoeologs exhibited the greatest reduction in fungal biomass. These results demonstrate the potential of prime editing to directly introduce resistance-associated alleles into wheat and provide a strategy for accelerating the development of rust-resistant cultivars.

## Results

Wheat rust diseases are persistent threats to wheat production because pathogens evolve rapidly. Adult-plant resistance (APR) genes often provide broad-spectrum, partial resistance and are considered more durable. The *Lr67* gene is an APR which confers broad-spectrum resistance to multiple biotrophic fungal pathogens, including leaf rust, stripe rust, stem rust, and powdery mildew <u>(Moore et al., 2015)</u>. *Lr67* encodes a D-genome Sugar Transport Protein 13 (*TaSTP13-D1*), which differs from the susceptible allele by two amino acid substitutions: G144R and V387L. The G144R substitution (TaSTP13^G144R^), encoded by a G-to-C transversion in *TaSTP13-D1* is sufficient to confer the resistance associated with *Lr67* <u>(Milne et</u> <u>al., 2024)</u>. Introgressing *Lr67* into elite bread wheat varieties requires extensive backcrossing, and the location on the D-genome renders it inaccessible to durum wheat breeding. For both bread wheat and durum wheat, the ability to directly introduce the functional *Lr67* allele through precision genome editing could accelerate deployment of this resistance.

Here, we used prime editing to convert a susceptible allele into a resistance-conferring variant by installing a single nucleotide substitution. Although early prime editing systems showed limited efficiency in wheat, the enhanced editor ePPEplus significantly improved editing efficiency in wheat protoplasts and regenerated plants <u>(Ni et al., 2023)</u>. Here, we adapted the ePPEplus editor to install the functional *Lr67* allele, TaSTP13^G144R^, into all three homoeologs of *TaSTP13* in hexaploid bread wheat. We also developed a wheat-compatible prime editing vector with high editing efficiency and simplified cloning for future trait engineering.

The ePPEplus system was adapted for *Agrobacterium*-mediated transformation in wheat by assembling the ePPEplus prime editor into a wheat-compatible acceptor vector JWPE1, which contains the *hygBr* hygromycin selection marker and the morphoregulators *GRF4-GIF1* to enhance regeneration (**Figure 1a; Supplemental Methods**). JWPE1 was adapted from the pJD633 CRISPR/Cas9 wheat transformation vector <u>(Debernardi et al., 2020)</u>. Expression of ePPEplus and engineered pegRNAs (epegRNAs) was driven by *ZmUbi* and *TaU6* promoters, respectively.

**Figure 1.**
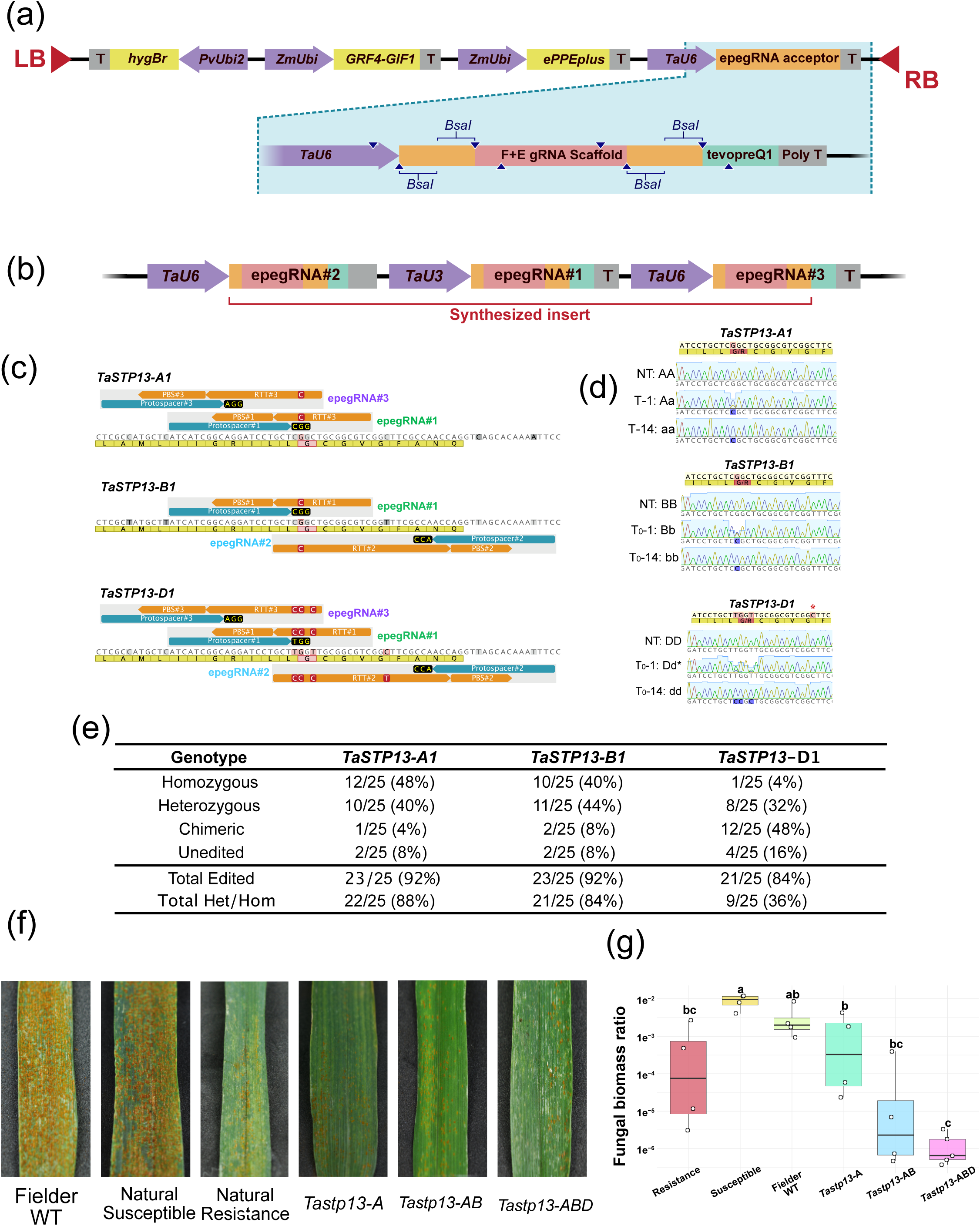
Prime editing of TaSTP13 homoeologs and assessment of leaf rust resistance. (a) Schematic of the wheat-compatible JWPE1 prime-editing vector containing *hygBr*, GRF4-GIF1 morphoregulators, and the ePPEplus editor. (b) Target regions of TaSTP13-A1, -B1 and -D1 showing protospacers, RTT/PBS designs and introduced edits. Additional synonymous substitutions were included in TaSTP13-D1 to introduce the MspA1I site. PAMs are shown in yellow; red boxes indicate the target nucleotide and G144; grey boxes indicate homoeolog-specific polymorphisms. (c) Representative Sanger chromatograms showing unedited, heterozygous, chimeric and homozygous edits. The expected but absent C-to-T substitution from epegRNA#2 is indicated by an asterisk. (d) TaSTP13^G144R^ editing frequencies in T0 plants. Total edited includes chimeric, heterozygous and homozygous events; Total Het/Hom includes heterozygous and homozygous events. (e) Leaf rust symptoms in control and prime-edited lines following *Puccinia triticina* inoculation. (f) Relative fungal biomass quantified by qPCR using PtITS normalized to TaEF-1α. Different letters indicate significant differences among genotypes (Tukey’s HSD, *n* = 4–5 biological replicates).

Three epegRNAs were designed to install the G-to-C transversion corresponding to the G144R amino acid substitution within the *TaSTP13* homoeologs (**Table S1**). A synthesized fragment encoding the three epegRNA cassettes was inserted into JWPE1 using BsaI Golden Gate cloning (**Figure S1, Sequence S1**). The three epegRNAs were expressed using alternating *TaU6* and *TaU3* Pol III promoters, which are more efficient for wheat prime editing than polycistronic tRNA processing systems <u>(Ni et al., 2023)</u>. Due to sequence divergence between the *TaSTP13* homoeologs, not all epegRNAs targeted all loci (**Figure 1b**). All three epegRNAs targeted *TaSTP13-D1*; epegRNA#1 and epegRNA#2 targeted *TaSTP13-B1*; and epegRNA#1 and epegRNA#3 targeted *TaSTP13-A1*. epegRNA#1 and epegRNA#2 were designed as a paired epegRNA strategy reported to improve wheat prime editing <u>(Lin et al., 2021)</u>. Reverse Transcription Template (RTT)s also introduced an *MspA1I* site to enable CAPS detection. In *TaSTP13-A1* and *TaSTP13-B1*, the G-to-C substitution alone creates the MspA1I site; however, due to sequence variation in *TaSTP13-D1*, two additional synonymous T-to-C substitutions in the RTT introduce this site. The resulting construct was introduced into the wheat cultivar Fielder lacking *Lr67* resistance allele, via *Agrobacterium*-mediated transformation <u>(Hayta et al., 2021)</u>.

Among the 31 transgenic plantlets, the MspA1I site was detected in 90.3%, 87.1% and 51.6% of lines at *TaSTP13-A1*, *-B1* and *-D1*, respectively (**Figure S2 and S3; Table S2**). Twenty-five of these plantlets regenerated into T_0_ plants (**Figure S4**) and were further analyzed by Sanger sequencing (**Figure 1c**). This revealed high editing efficiencies for the TaSTP13^G144R^ allele: 92.0% in *TaSTP13-A1*, 92.0% in *TaSTP13-B1*, and 84.0% in *TaSTP13-D1* (**Figure 1d; Table S3**). These efficiencies included chimeric, heterozygous, and homozygous editing events. Homozygous TaSTP13^G144R^ edits were observed in 48.0% of T_0_ plants in *TaSTP13-A1*, 40.0% in *TaSTP13-B1*, and 4.0% in *TaSTP13-D1*. One line (T_0_-14) was homozygous for the edited allele in all three genomes (**Figure 1d**). No byproducts were observed in any of the Sanger chromatograms (**Figure S5**). The lower frequency of homozygous D-genome edits may reflect the requirement for multiple nucleotide changes at this locus. The RTT design predicted an additional synonymous C-to-T substitution at TaSTP13-D1 when using epegRNA #2, but this change was not detected (**Figure S5**). The RTT design predicted an additional synonymous C-to-T substitution at *TaSTP13-D1* when using epegRNA #2, but this change was not detected (Figure S5), suggesting that epegRNA#2 did not contribute to editing at *TaSTP13-D1*, potentially because of its very long RTT.

To assess whether stacking homoeolog edits enhanced resistance, we evaluated a stepwise series of *TaSTP13-A*, *TaSTP13-AB* (AB), and *TaSTP13-ABD* (ABD) edited genotypes. T_0_ plants were advanced to T_3_ to reduce contributions from mosaic or chimeric events and evaluate stable inheritance (**Table S4** and **Figure S6**). The genotypes evaluated included the susceptible control (Fielder WT), a naturally susceptible line and its sister line carrying the resistance-associated *TaSTP13-D1* allele (resistant line), as well as the prime-edited line. Disease severity was evaluated using the CIMMYT rust scoring scale (**Table S5**). Fielder WT and the susceptible line developed extensive pustule formation, whereas the resistant control showed minimal infection (**Figure 1e**). Among the prime-edited lines, *TaSTP13-A* exhibited visible pathogen colonization, but reduced disease severity compared with the susceptible controls. The *TaSTP13-AB* and *TaSTP13-ABD* lines also displayed reduced disease severity relative to both susceptible genotypes (**Figure 1f**). To validate these observations, fungal biomass was quantified by qPCR using wheat *TaEF-1α* and fungal *PtITS* markers (**Figure S7**). The susceptible line had the highest fungal biomass, followed by Fielder WT. Edited lines had lower fungal biomass than susceptible controls, while no significant differences were observed between the resistant control and edited lines (**Figure 1f**). *TaSTP13-AB* showed an intermediate phenotype, whereas *TaSTP13-ABD* had the greatest significant reduction in fungal biomass and the strongest difference from susceptible controls. We also examined the expression of genes associated with plant defense responses by monitoring the expression of Pathogenesis-related protein -1 (PR1), -3 (PR3), and superoxide dismutase (SOD) over a time course following inoculation. PR1 and PR3 showed upregulation at 12 and 24 h post-inoculation (hpi), with elevated expression maintained at 48 hpi. In contrast, SOD expression showed a pronounced downregulation at 48 hpi. Edited lines showed stronger expression changes than susceptible controls (**Figure S8**), consistent with an enhanced or more rapidly activated defense response. Together, disease scoring, fungal biomass, and defense-gene expression support enhanced leaf rust resistance following installation of *TaSTP13^G144R^* allele via prime editing.

Our study demonstrates the first installation of a functional disease-resistance allele in wheat by gene editing. Using prime editing, we installed *TaSTP13^G144R^* across all three *TaSTP13* homoeologs in hexaploid bread wheat. Using the JWPE1 vector containing *GRF4-GIF1* morphoregulators <u>(Debernardi et al., 2020)</u>, we achieved high editing frequencies in regenerated T_0_ plants (84.0–92.0%). This work established prime editing as a potential approach for introducing a resistance-associated allele into wheat without linkage drag. Our resistance assays were conducted under controlled inoculation conditions, however, field performance, durability across pathogen populations, and agronomic effects remain to be determined. To facilitate further investigation and potential application of this approach in wheat improvement, the prime-editing construct (**Figure S9**) developed in this study will be made publicly available through Addgene.

## Acknowledgement

We thank Dr. Sridhar Bhavani, Dr. Naeela Qureshi and Dr. Pawan Singh for their support with the rust assays, as well as the technical staff Hedilberto Velazquez. We also acknowledge Ruben Basilio and Ivan Martínez for their contributions to plant maintenance and establishment of *Agrobacterium*-mediated transformation. We thank Dr. Kevin Pixley for prioritization of this work and help in acquisition of the funding.

## Supplemental Information

### A) Supplemental Methods

#### JWPE1 plasmid construction

To develop the ePPEplus wheat-compatible expression vector (named JWPE1), the JD633 CRISPR/Cas9 wheat transformation vector (Addgene, Plasmid #160393, https://www.addgene.org/160393/) <u>(Debernardi</u> <u>et al. 2020)</u> was adapted to express ePPEplus and engineered pegRNAs (epegRNAs) as described by Ni et al. (2023). The Cas9 coding sequence and guide RNA expression cassette were excised from pJD633 using SbfI and PmeI (New England Biolabs, Ipswich, Massachusetts, US) and replaced with a synthesized fragment containing the codon-optimized ePPEplus coding sequence (modified to remove an internal SbfI site) and an epegRNA expression cassette with compatible overhangs (GenScript, Piscataway, New Jersey, US) (**Sequence S2**). The *TaU6* promoter present in pJD633 was retained to drive epegRNA expression. The epegRNA scaffold incorporated the ‘flip and extension’ (F+E) guide RNA scaffold <u>(Chen</u> <u>et al. 2014; Ni et al. 2023)</u>. **Sequence S3** is the final ePPEplus expression cassette sequence, and **Sequence S4** is the final epegRNA expression cassette sequence. A labeled plasmid map of JWPE1 is found in **Figure S9**.

#### epegRNA design and assembly into JWPE1

Three epegRNAs were designed to install the G-to-C substitution corresponding to the G144R amino acid substitution in the *TaSTP13* homoeologs. Protospacer, RTT, and PBS sequences were designed using PlantPegDesigner (http://www.plantgenomeediting.net/) <u>(Jin et al. 2023)</u>. A linker sequence was added between the PBS and tevopreQ1 motif using pegLIT (https://peglit.liugroup.us/) <u>(Nelson et al. 2022)</u>. The three epegRNA cassettes were synthesized as a single fragment (GenScript, Piscataway, New Jersey, US) (**Sequence S1**) with flanking BsaI overhangs and cloned into JWPE1 using Golden Gate assembly with BsaI (New England Biolabs, Ipswich, Massachusetts, US), producing JWPE1-STP13^G144R^. Table S1 provides detailed sequences for the protospacer, RTT, PBS, and linker of each epegRNA.

#### *Agrobacterium*-mediated transformation of wheat

Wheat transformation was carried out in the cultivar Fielder following the protocol of <u>Hayta et al. (2021)</u>. The JWPE1-STP13^G144R^ construct containing all three epegRNAs was introduced via *Agrobacterium tumefaciens* strain EHA105. Transformed calli were selected using hygromycin, and regenerated plantlets were transferred to soil in glasshouses. Transgenic plants were confirmed by PCR analysis using primers specific to the *hygBr* transgene (**Table S6**).

#### CAPS genotyping

Genomic DNA of each plant was extracted from the leaves using the CTAB DNA extraction method. Homoeologs-specific PCR primers were designed to amplify a 783 bp region for *TaSTP13-A1*, a 1000 bp region for *TaSTP13-B1*, and a 1,203 bp region for *TaSTP13-D1*. After PCR with Phusion High-Fidelity DNA Polymerase (New England Biolabs, Ipswich, Massachusetts, US), amplicons were purified using a GeneJET PCR Purification Kit (Thermo Fisher, Waltham, Massachusetts, US). A synthetic positive control fragment (+C) containing the G-to-C substitution and *MspA1I* site was created for *TaSTP13-A1* using primer extension mutagenesis. PCR products were digested using *MspA1I* (New England Biolabs, Ipswich, Massachusetts, US) at 37°C for 2 hours and resolved on a 2% agarose gel. Edited alleles introduced an *MspA1I* site, producing 465 bp and 318 bp fragments for *TaSTP13-A1*; 508 bp, 449 bp, and 43 bp fragments for *TaSTP13-B1* (an *MspA1I* site in the unedited fragment produces 957 bp and 43 bp fragments); 697 bp and 506 bp fragments for *TaSTP13-D1* (**Figure S3a**). Primer sequences for amplification of *TaSTP13* homoeologs and primer extension mutagenesis are found in **Table S6**.

#### PCR amplification, Sanger sequencing, and editing outcome analysis

Genomic regions of each *TaSTP13* homoeologs were amplified using specific primers (**Table S6**). PCR products were purified directly using the GeneJET PCR Purification Kit (Thermo Fisher Scientific), following the manufacturer’s instructions, with a final elution volume of 20 µL. Purified PCR products were submitted to Psomagen for Sanger sequencing using the ABI 3730xl DNA Analyzer (Applied Biosystems. Sequence chromatograms were analyzed using TIDER (Tracking of Insertion, DEletions and Recombination events; https://apps.datacurators.nl/tider/), a web-based tool for quantifying template-directed genome-editing outcomes by sequence-trace decomposition <u>(Brinkman et al. 2018)</u>. TIDER compares the chromatogram of the edited sample with those of an unedited control and a reference sequence containing the expected edited nucleotide(s). The analysis was used to assess the editing outcomes at each *TaSTP13* homoeolog.

#### Plant growth conditions and leaf rust assay

Seeds of the susceptible control (Fielder WT), susceptible (RL6077/AOC YR -), sister resistant line (RL6077/AOC YR +), and prime edited *Lr67* lines were grown in individual pots (Maceta No. 7; https://plasticosalica.com.mx/) containing a substrate mixture of 40% steam-sterilized PittMoss and 60% field soil at the CIMMYT Biosafety Laboratory. Plants at adult stage were inoculated with *Puccinia triticina* strain MJB/SP urediniospores resuspended in mineral oil (Soltrol® 220 Isoparafina) using a pump sprayer (Bomba de Alto Vacío FE-1376 300 l/min, https://felisa.com.mx/) Following inoculation, plants were allowed to dry for 15 min and then transferred to a custom-made plastic chamber equipped with a humidifier (Vitallys Plus VUH-3) to maintain high relative humidity. Plants were incubated in the chamber for 16 h under dark conditions. Subsequently, plants were maintained under controlled environmental conditions with a 16:8 hrs (light:dark photoperiod), day/night temperatures of 24 °C and 22 °C, respectively, and watered three times per week with municipal water. Fertilization was applied three times prior to the rust assay using YaraMila (https://www.yara.us/) 21-17-3 (NPK). At the heading stage (Zadoks growth stage 55–59), plants were transferred to a CMP6050 Conviron (https://www.conviron.com/) growth chamber set to 85% relative humidity, 16 h light/8 h dark photoperiod and temperature of 23 °C and 20 °C during the light/dark cycles. To promote infection, two misting devices were installed to provide humidity prior to the onset of the light cycle (**Supplementary Fig. S10**). A mock treatment was included under the same experimental conditions, with plants inoculated with mineral oil only. Disease severity was evaluated according to the CIMMYT rust scoring protocol (http://hdl.handle.net/10883/1109).

#### Quantification of *Puccinia triticina* DNA by qPCR

Leaf samples from the genotypes used in the rust assay were collected at 10 days post inoculation and freeze-dried prior to genomic DNA extraction using the CTAB method <u>(Dreisigacker et al. 2013)</u>. Total DNA was used for quantitative PCR (qPCR) using the Luna Universal qPCR & RT-qPCR Kit (New England Biolabs), following the manufacturer’s instructions. Standard curves were generated using genomic DNA from the genotype Fielder WT, with the housekeeping genes elongation factor 1-alpha (*TaEF1α*) used for normalization. Fungal biomass was quantified using *Puccinia triticina*-specific ITS primers. Amplification was performed on the 7500 Fast Real-Time PCR System. Data analysis was conducted in R (v4.5.1). All primers used for this study are in Table S6.

#### RNA extraction and RTqPCR analysis

Leaf samples from the genotypes used in the rust assay were collected at 0 (pre-inoculation), 12, 24, and 48 hpi, immediately frozen in liquid nitrogen, and stored at −80 °C until processing. Total RNA was extracted using the Quick-RNA Plant Miniprep Kit (Zymo Research), following the manufacturer’s instructions. Genomic DNA contamination was removed by DNase I treatment according to the manufacturer’s protocol of Invitrogen. cDNA synthesis was performed using the Transcriptor High Fidelity cDNA Synthesis Kit (Roche), following the manufacturer’s guidelines. RTqPCR was carried out on a 7500 Fast Real-Time PCR System (Applied Biosystems) using a 1× Luna Universal qPCR & RT-qPCR Kit (New England Biolabs). Each reaction contained 0.25 µL of each primer and a 1:20 (v/v) cDNA dilution, for a final volume of 10 µL. Thermal cycling conditions were as follows: initial denaturation at 95 °C for 60 s, followed by 40 cycles of denaturation at 95 °C for 15 s and annealing/extension at 60 °C for 60 s. A melting curve analysis was performed at the end of amplification to verify product specificity using the instrument software’s default settings. Mock samples were used to calculate the relative expression change based on the Double delta Ct analysis (2^-ΔΔCt^).

## B) Supplemental Figures

**Supplementary Figure 1.**
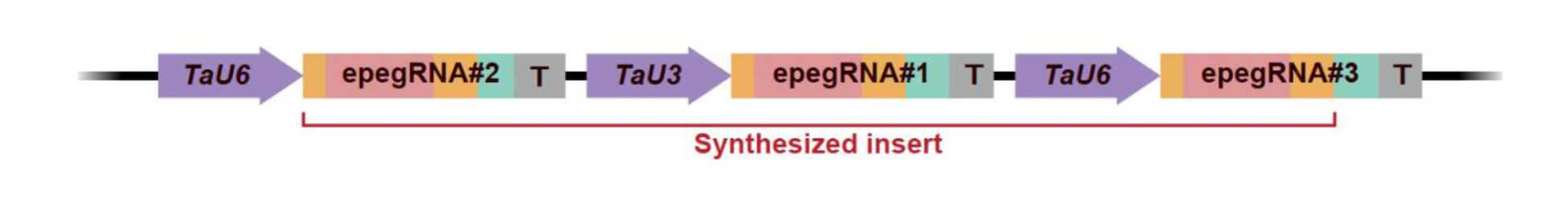
**epegRNA cassette used to install the TaSTP13^G144R^ substitution.**

**Supplemental Figure 2.**
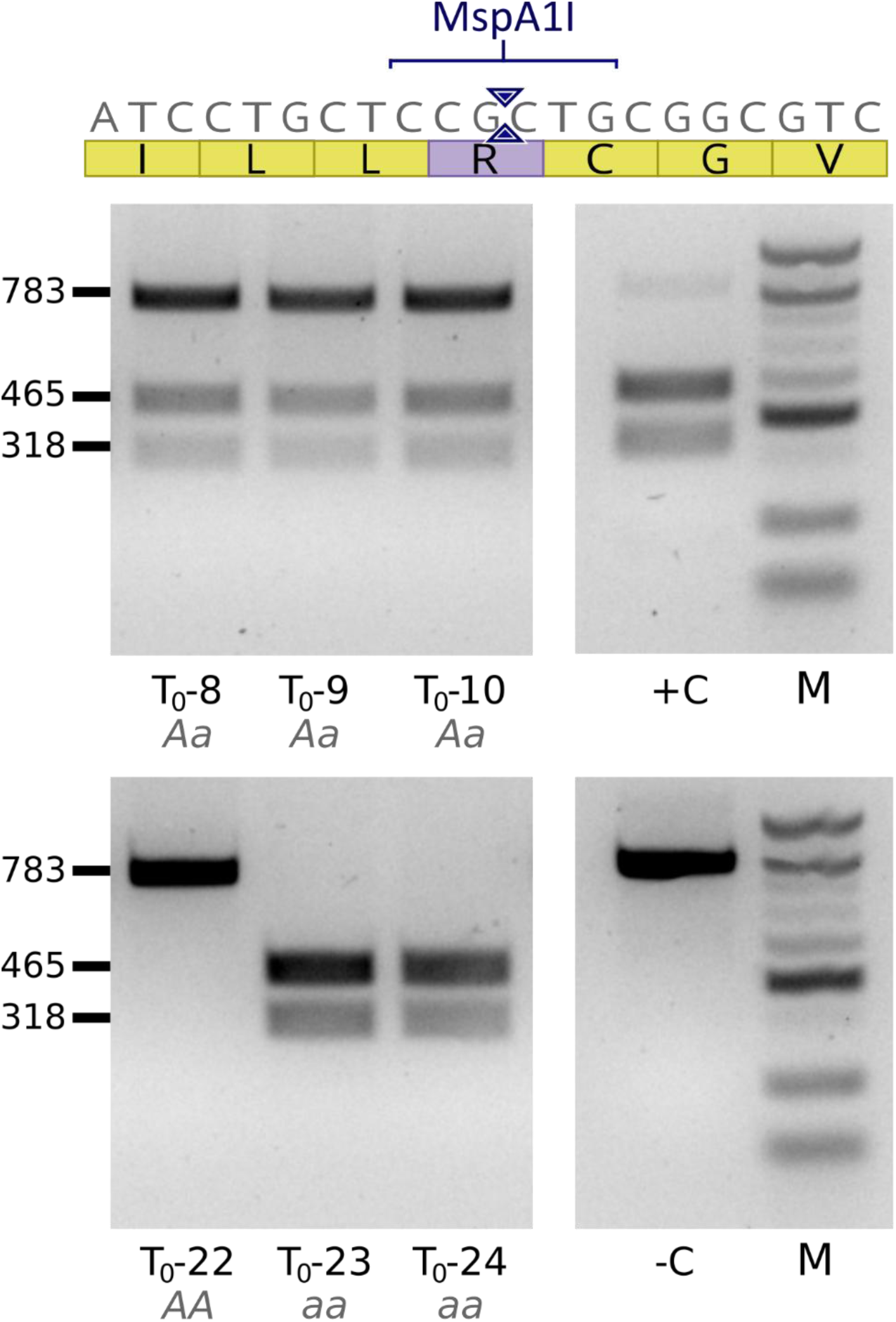
Representative CAPS genotyping of *TaSTP13-A1*. The G-to-C edit introduces an MspA1I restriction site that cleaves the 783 bp amplicon into 465 bp and 318 bp fragments. The pictures shown are homozygous (aa, fully digested), heterozygous (Aa, mixed), and wild-type (AA, uncut) lines alongside a synthetic positive control (^+^C), a non-transgenic negative control (^-^C), and a 100 bp DNA ladder (M).

**Supplemental Figure 3.**
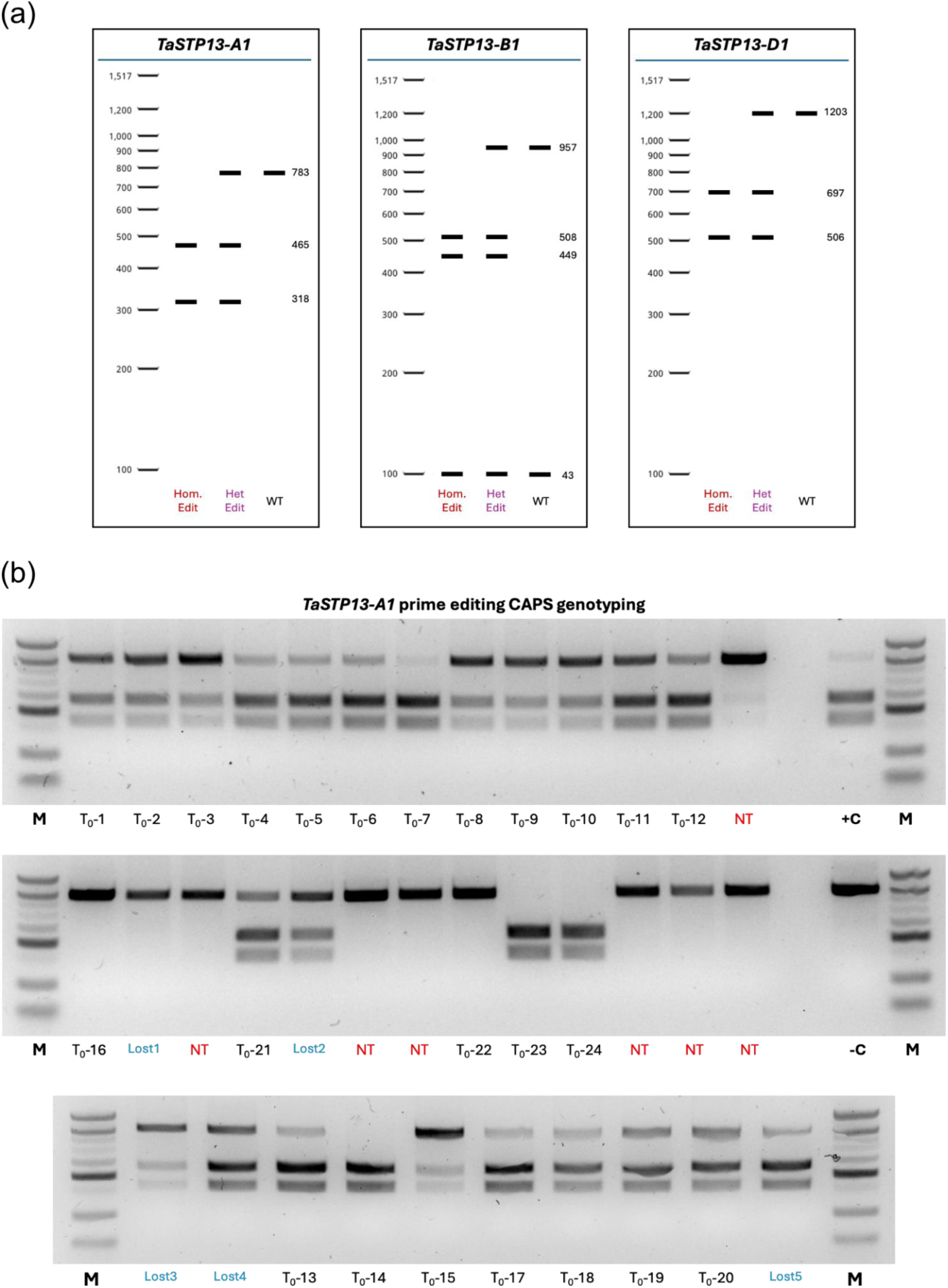
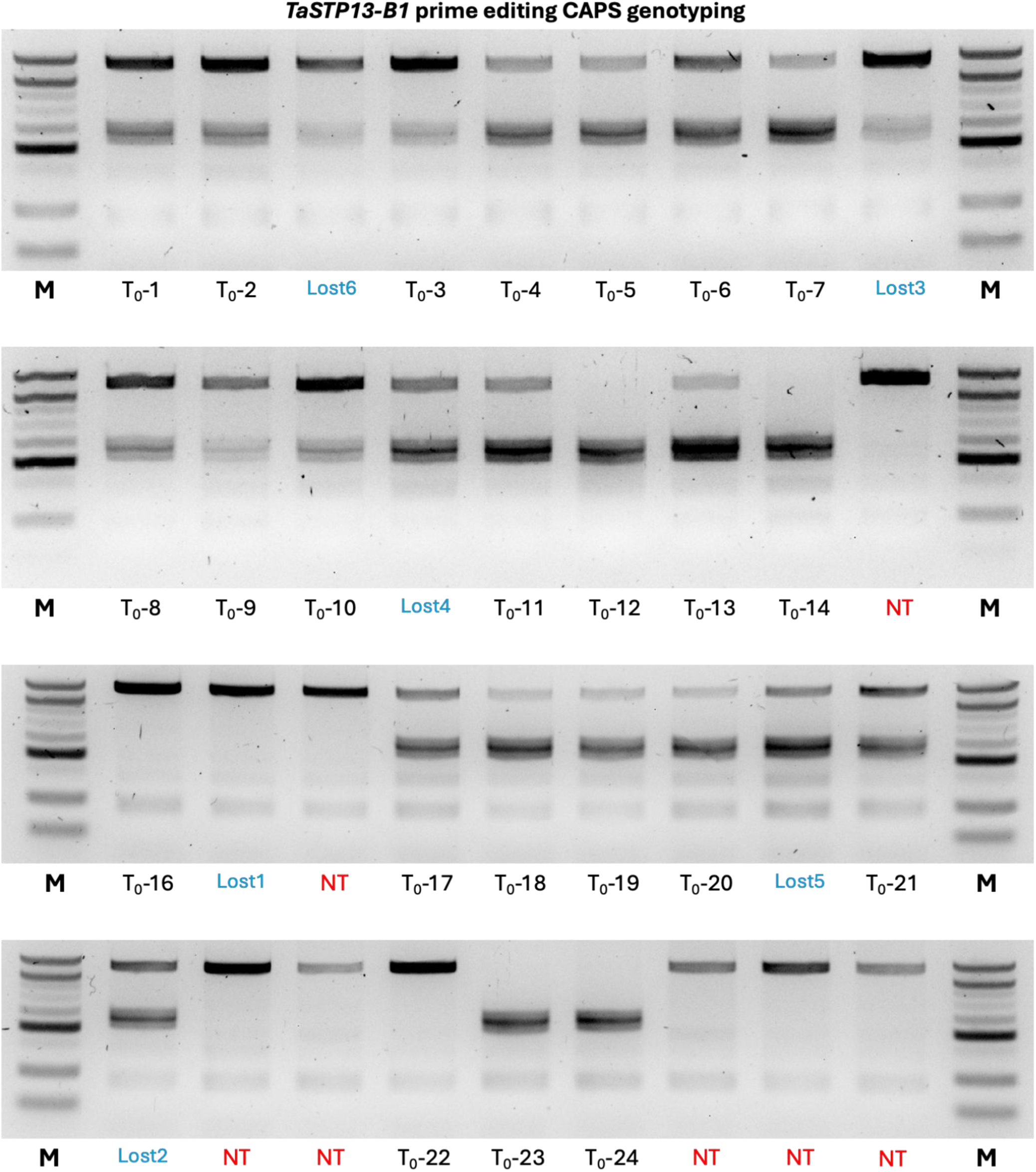
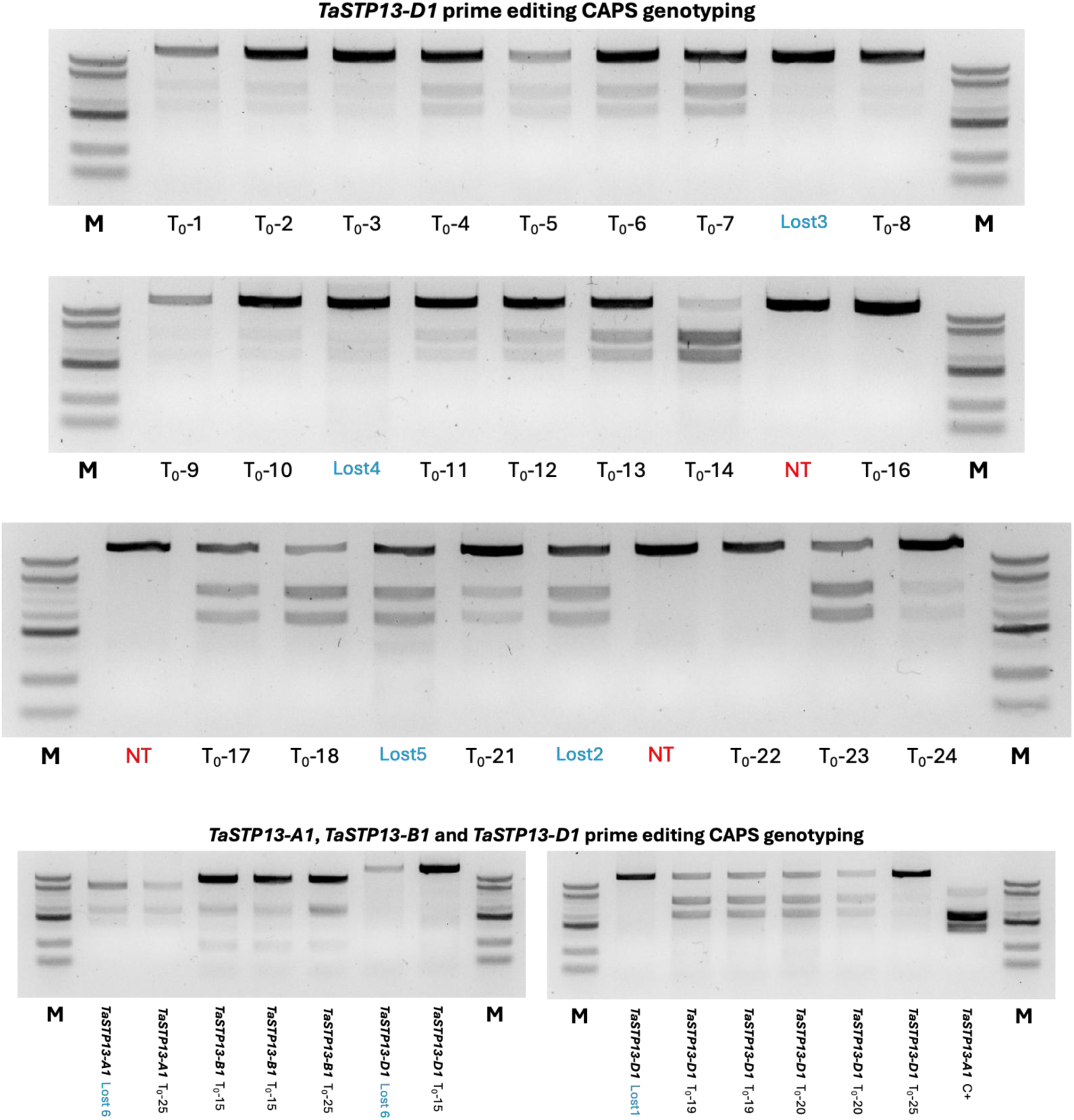
Complete CAPS genotyping gels for *TaSTP13-A1*, *-B1*, and *-D1* in T_0_ plantlets. The correct installation of the G144R substitution introduces an MspA1I restriction site used for CAPS detection. (a) Virtual gel showing expected banding patterns for unedited, heterozygous, and homozygous genotypes. (b) Gel electrophoresis images of CAPS-digested PCR products for all T_0_ lines. Lines labelled as “LostX” did not regenerate and were therefore excluded from Sanger sequencing analysis. NT: non-transgenic; -C: negative wild-type control; +C: synthetic *TaSTP13-A1* positive control; M: 100 bp DNA ladder.

**Supplemental Figure 4.**
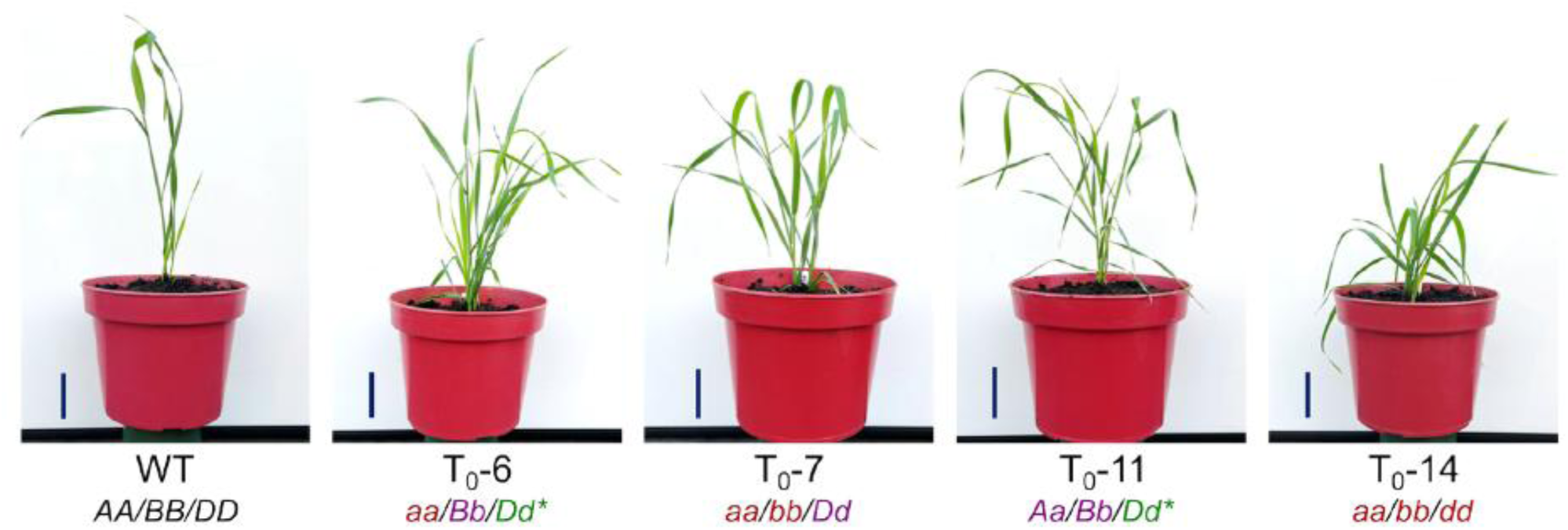
Representative T₀ wheat plants regenerated with the TaSTP13G144R allele installed. The Sanger sequencing genotype for each plant is shown below its corresponding plant ID. Scale bar = 5 cm. WT: Wild-type control; AA/BB/DD: unedited; Aa/Bb: heterozygous edit; Dd*: chimeric edit; aa/bb/dd: homozygous edit.

**Supplemental Figure 5.**
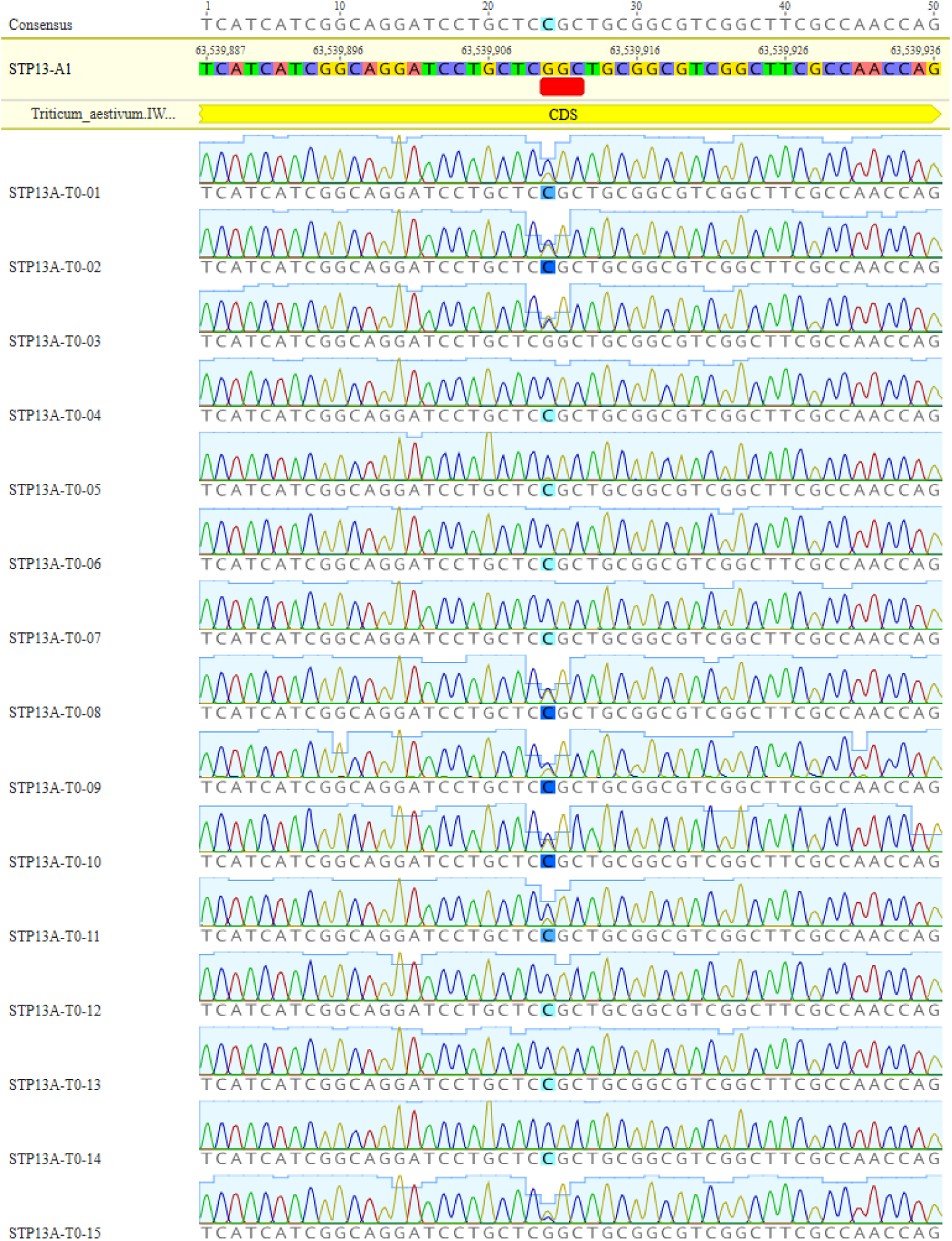
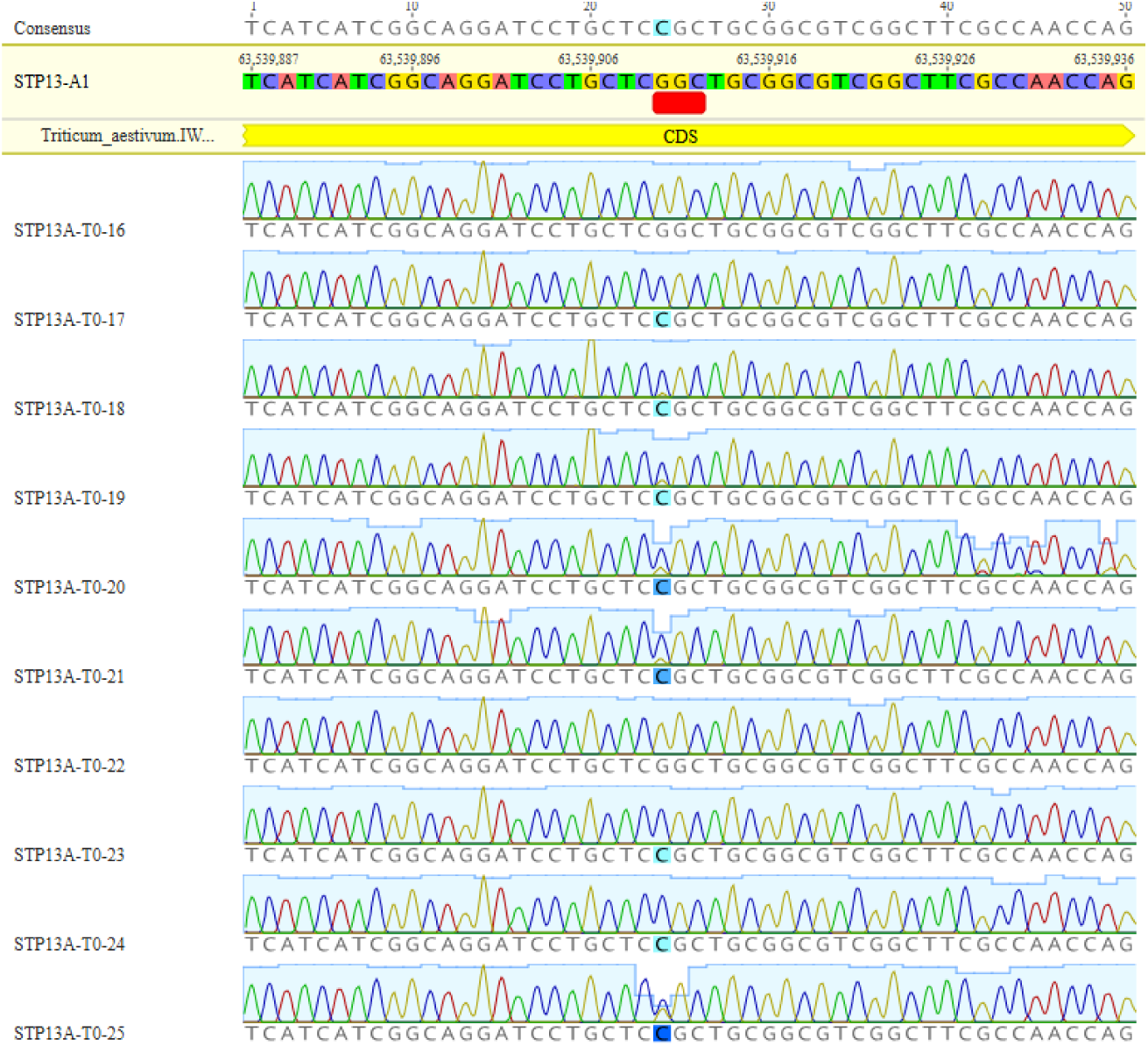
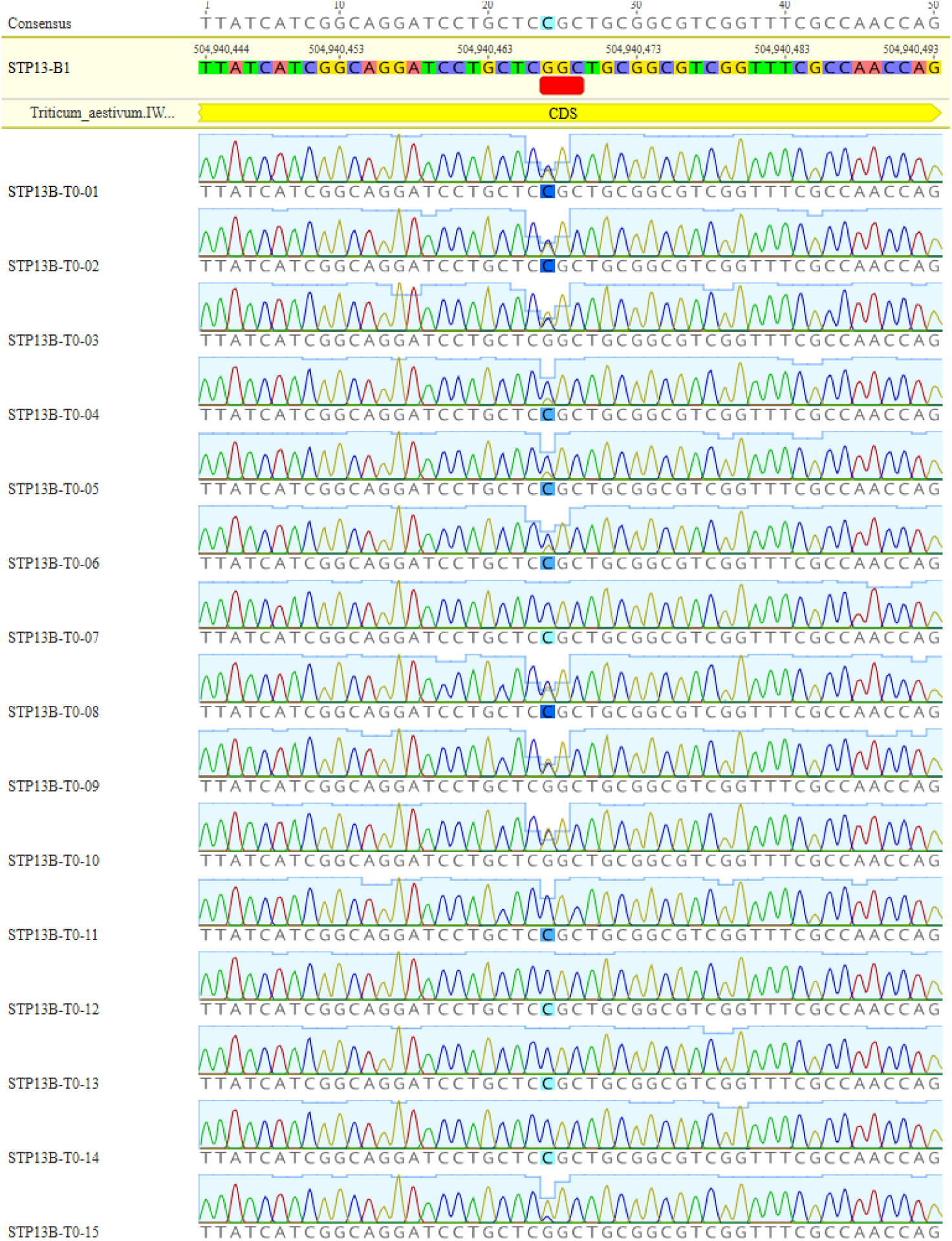
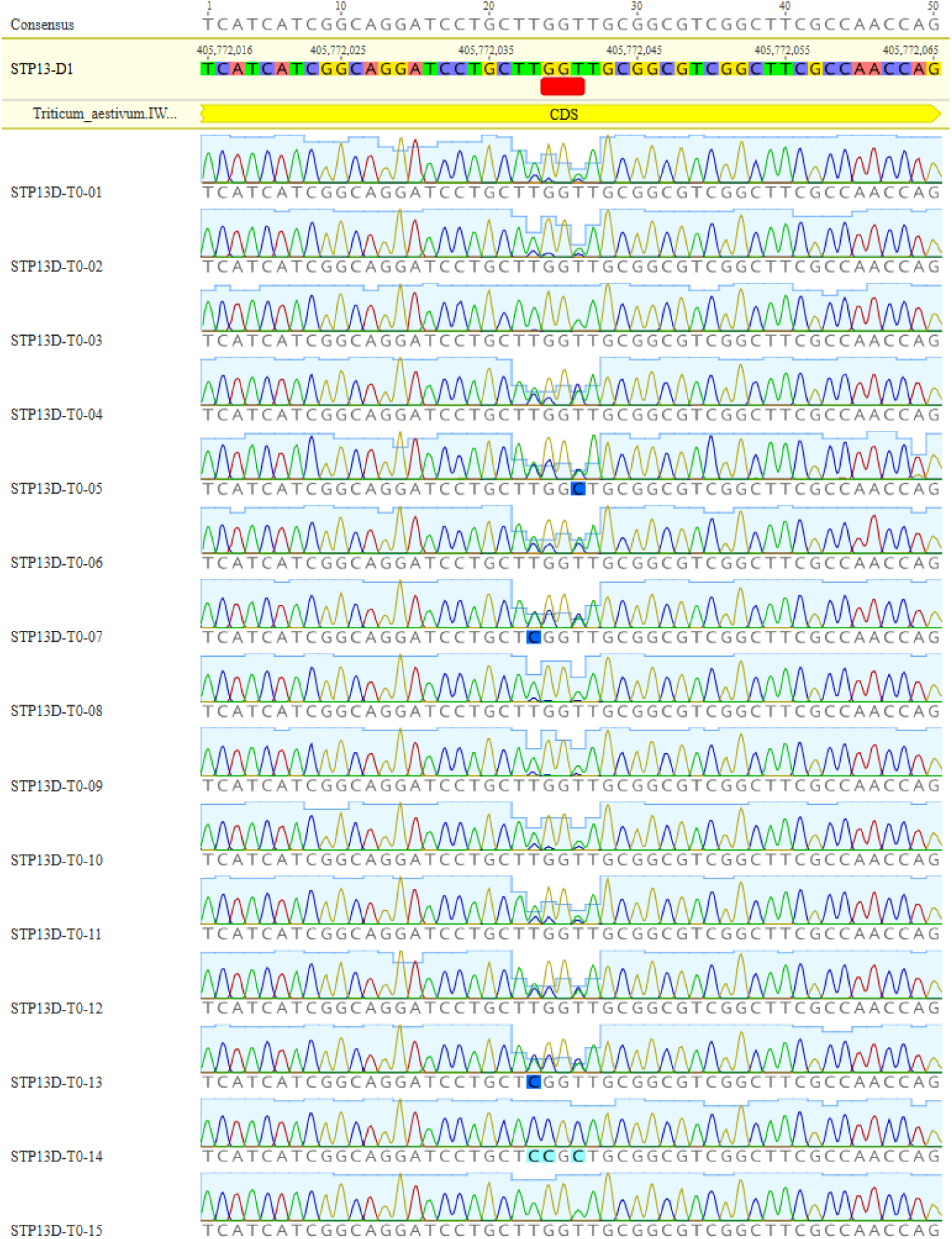
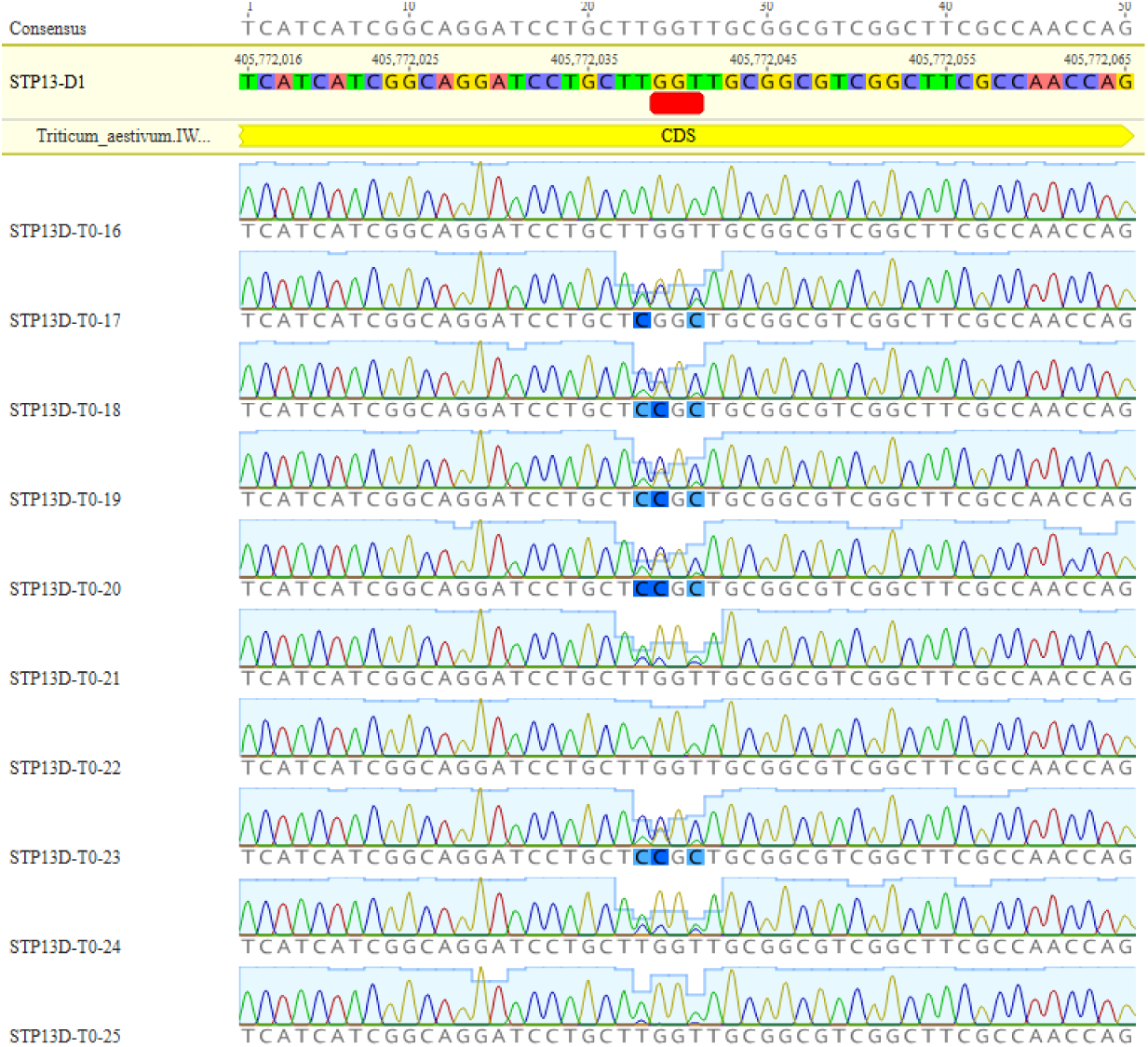
All Sanger sequencing chromatograms of T_0_ regenerated plants for *TaSTP13-A1*, *-B1*, and *-D1*. The red rectangle highlights the codon corresponding to G144. Cytosine base colors reflect quality scores, with dark blue indicating low confidence and light blue indicating high confidence.

**Supplemental Figure 6.**
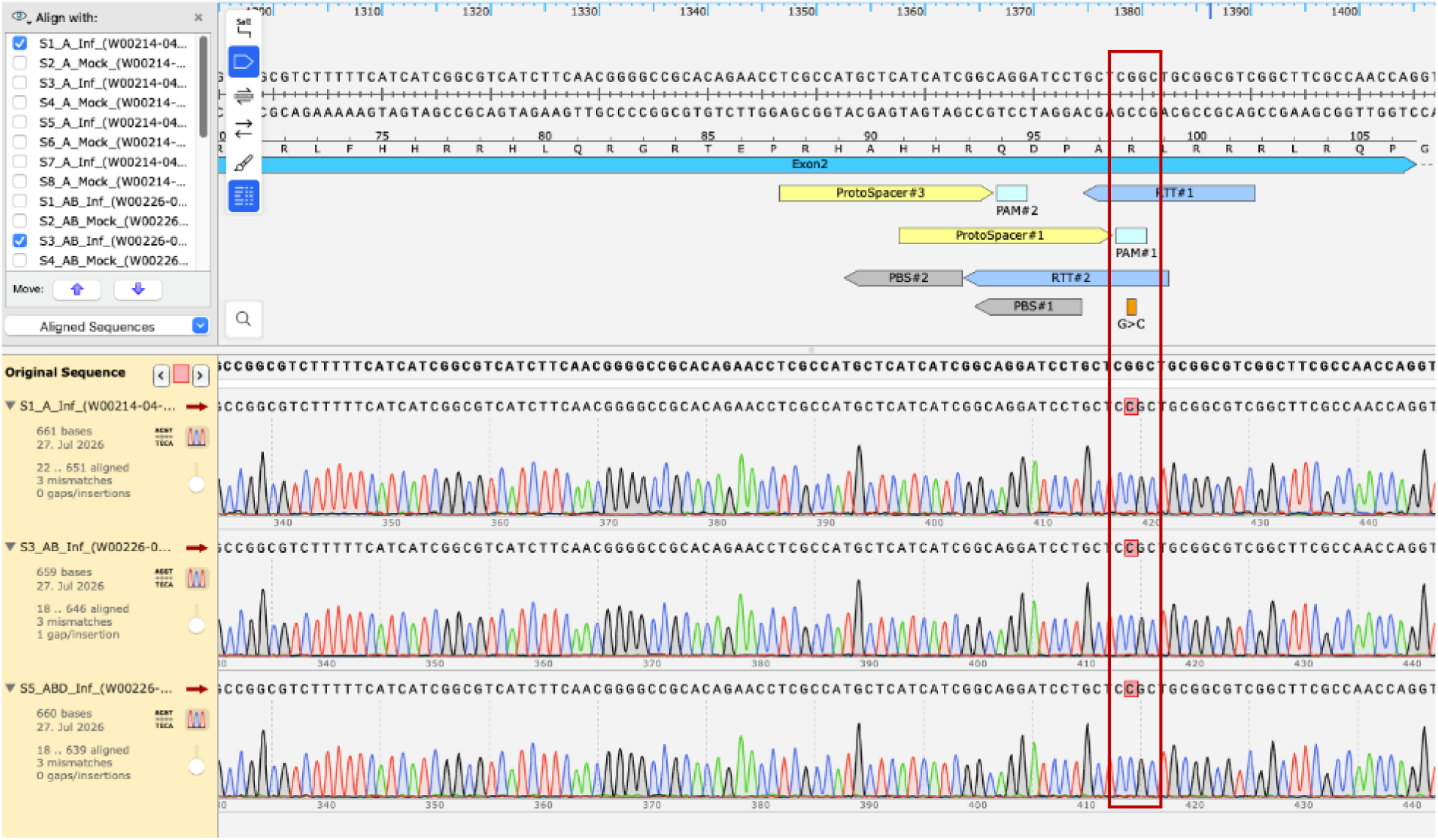
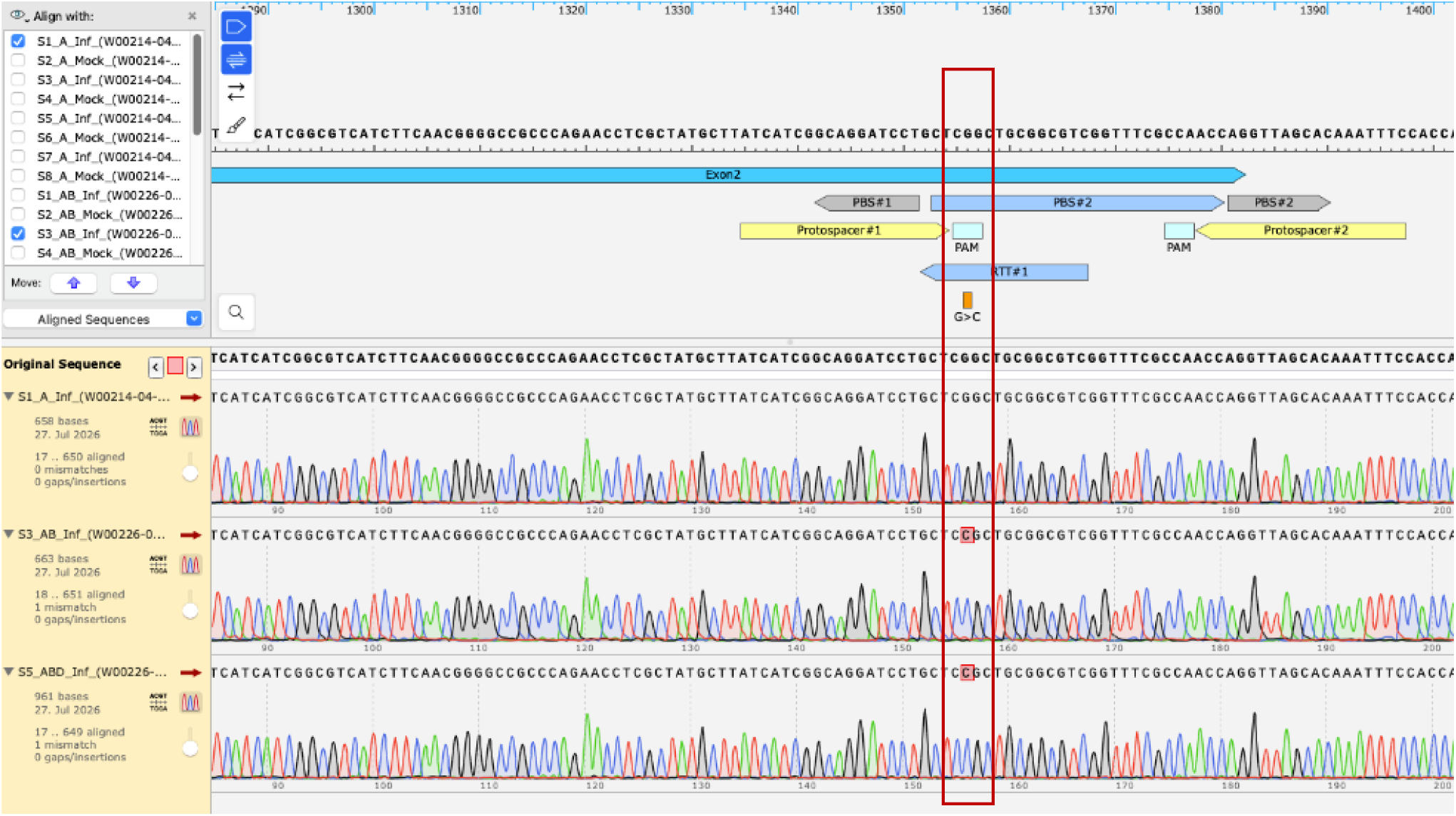
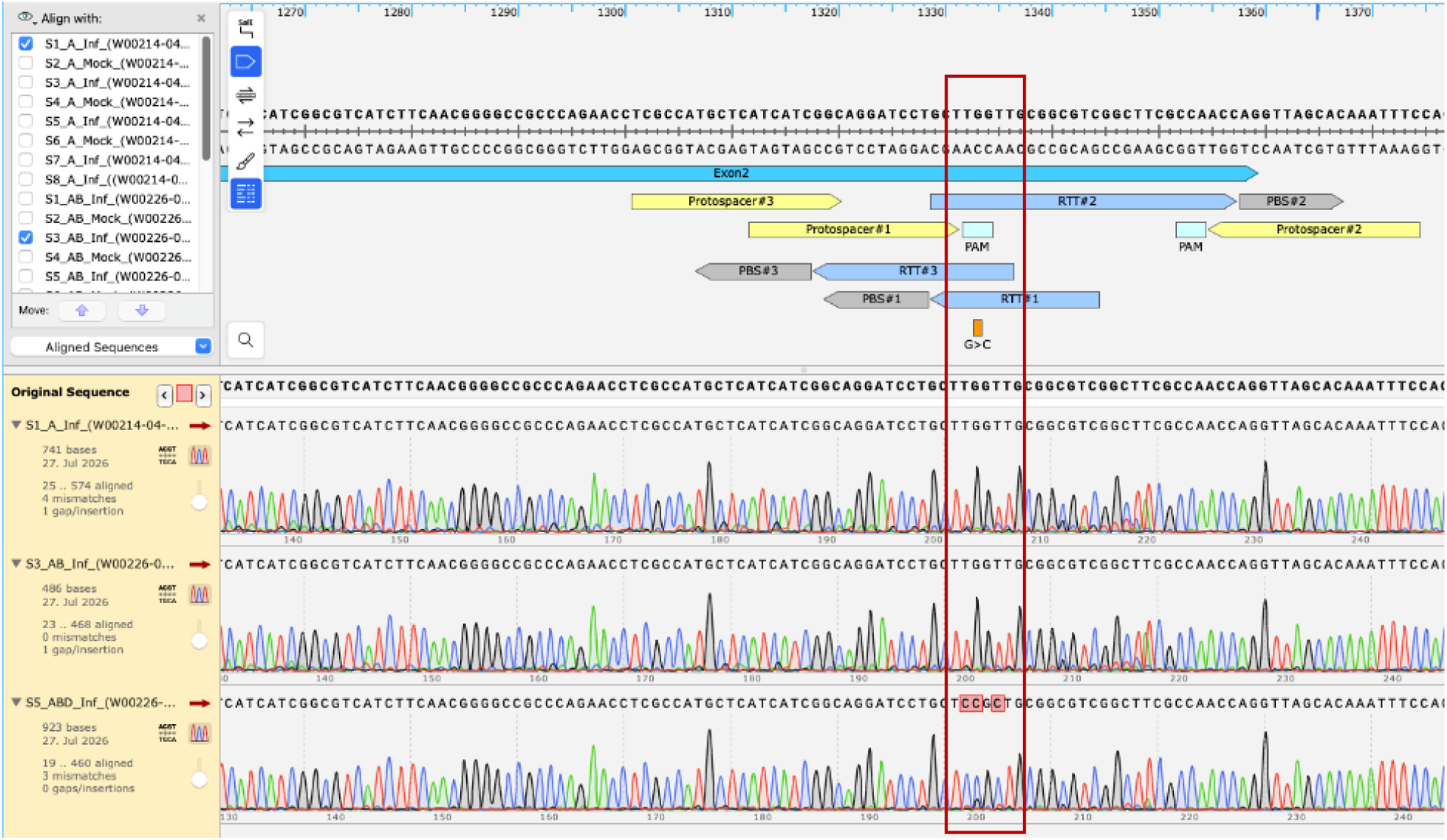
Representative T_3_ wheat chromatograms for *TaSTP13-A1*, *-B1*, and *-D1.* Samples shown are S1_A_Inf, S3_AB_Inf, and S5_ABD_Inf, representing all three wheat genomes. Red boxes indicate the nucleotide changes corresponding to each edited genome. All chromatograms are in **Sequence S5**.

**Supplemental Figure 7.**
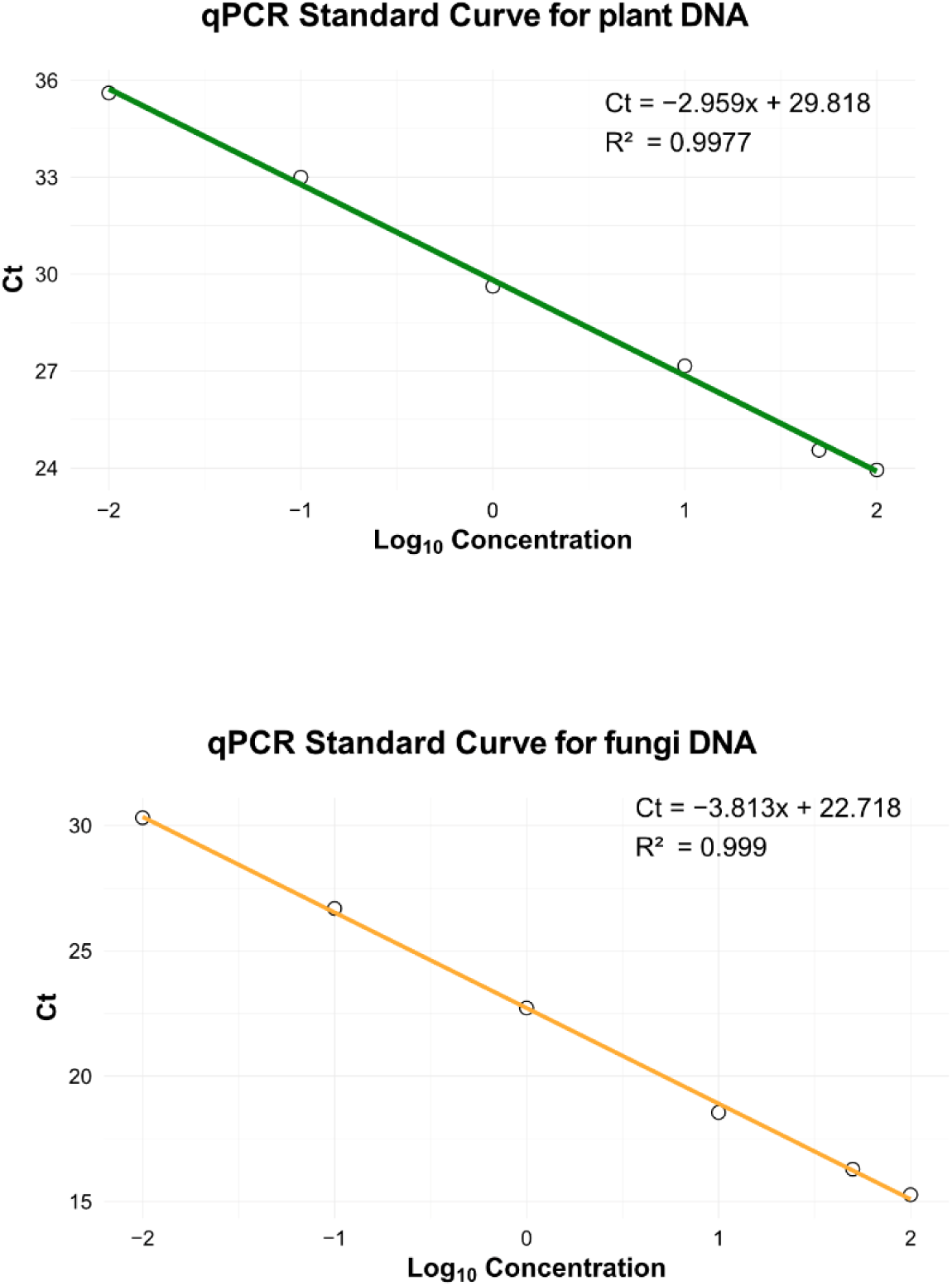
qPCR standard curves. Standard curves for wheat and fungal DNA quantification were generated using the wheat reference gene *TaEF-1α* and the fungal marker *PtITS*, respectively. DNA concentrations ranging from 100 to 0.01 ng per reaction (100, 50, 10, 1, 0.1, and 0.01 ng) were used to generate the standard curves.

**Supplemental Figure 8.**
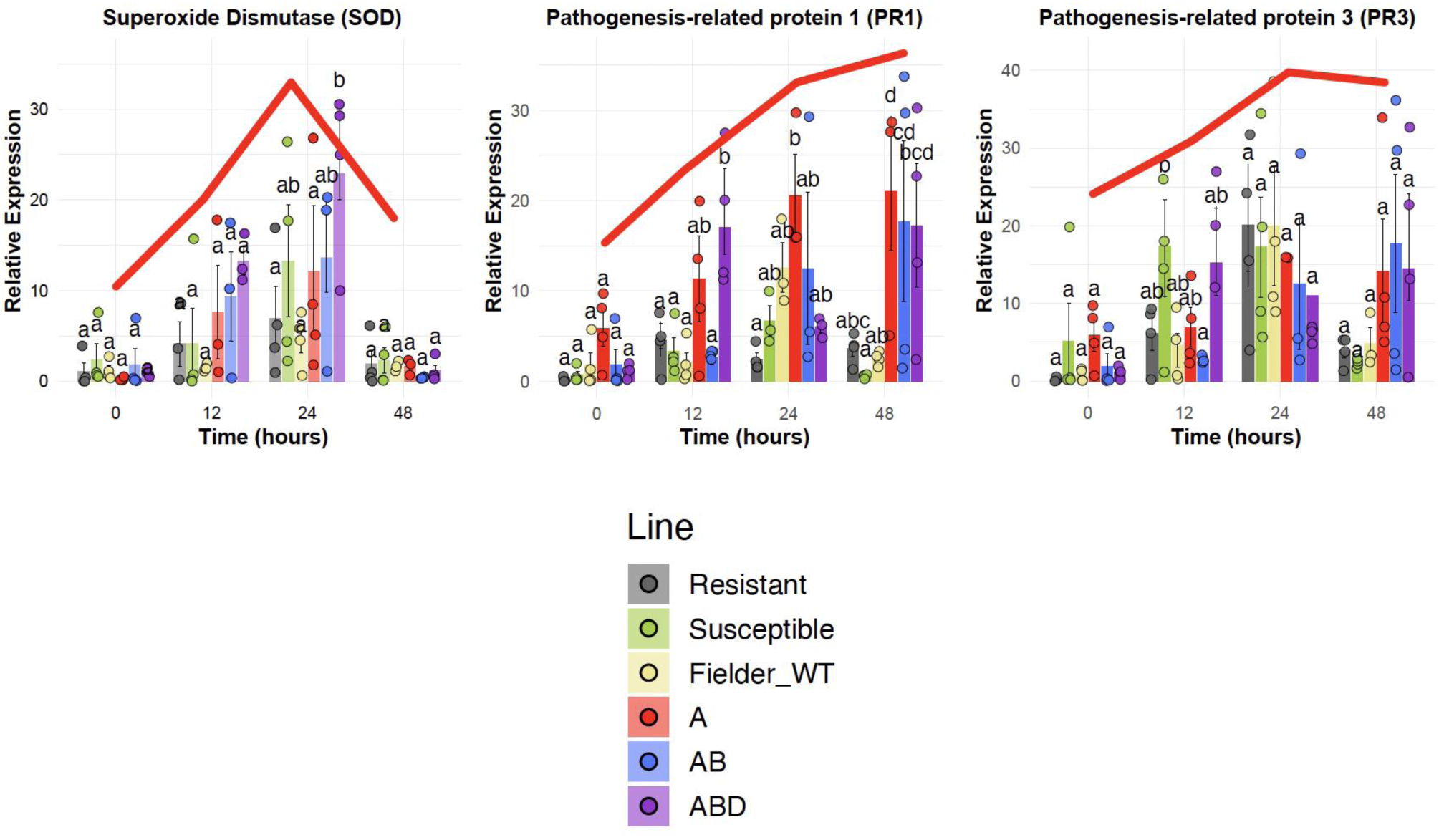
Expression analysis of three APR genes in control and edited lines at 0, 12, 24, and 48 h post-inoculation. Changes in transcript abundance for each gene were assessed at each time point relative to the respective control. The red line represents the overall temporal expression trend which was consistent with the expected expression pattern.

**Supplemental Figure 9.**
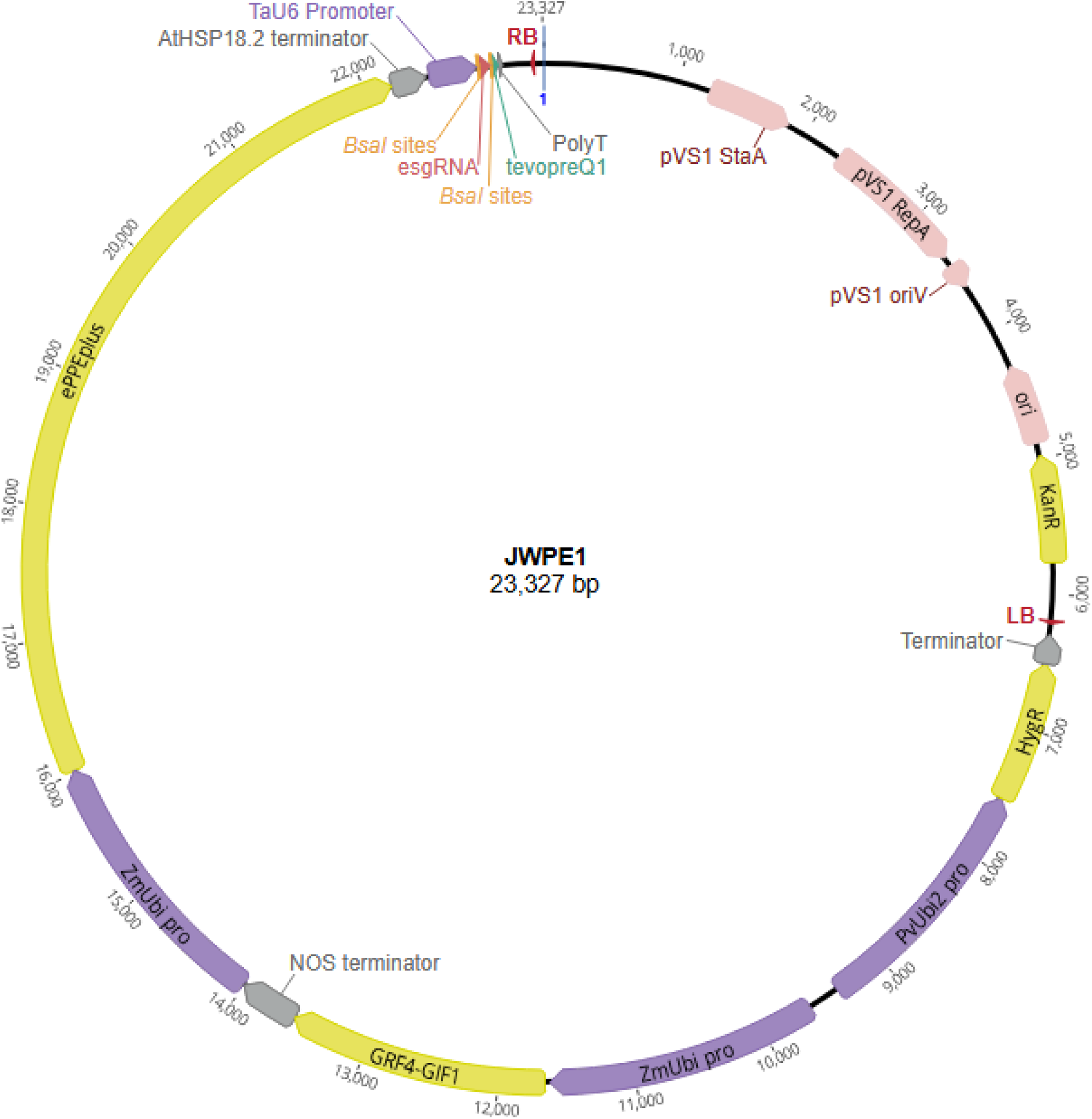
Labelled plasmid diagram of JWPE1. Promoters are indicated by purple, CDS are indicated by yellow, and terminators are indicated by grey. RB: right border, LB: left border, pro: promoter.

**Supplemental Figure 10.**
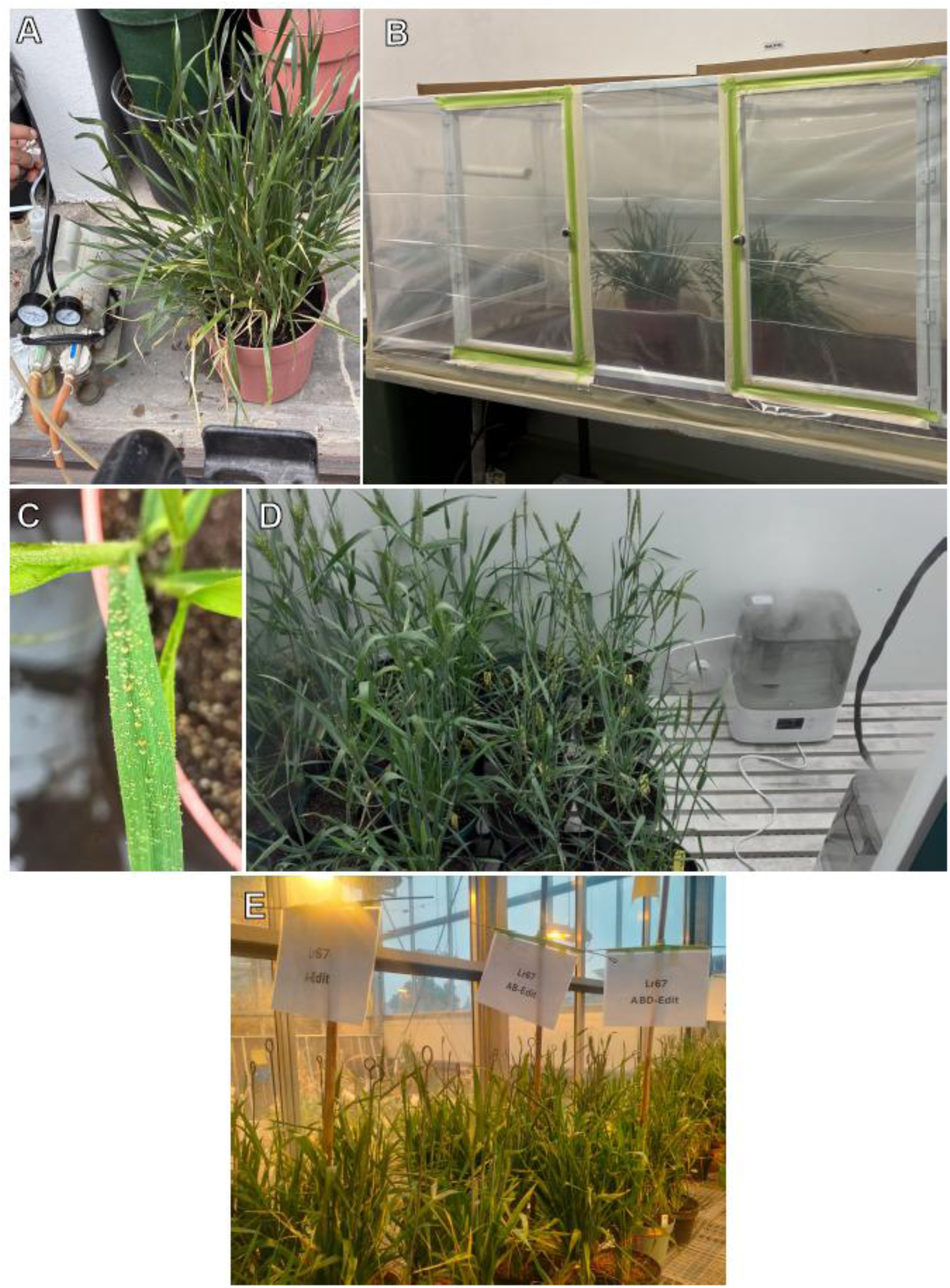
Rust assay experimental settings. A) Application of *P. triticina* urediniospores resuspended in mineral oil to wheat plants using a pump sprayer. B) Plants maintained for 16 hours under dark and high-humidity conditions in a custom-made plastic chamber equipped with a humidifier to promote urediniospore germination and infection. C) Rust infection activation on the leaf surface after exposure to moisture. D) Plants maintained under controlled environmental conditions in a Conviron growth chamber (CMP6050; 85% relative humidity, 16 h light/8 h dark photoperiod, and 23/20 °C day/night temperatures), with misting devices used to increase humidity before the onset of the light period. E) Mock-treated plants maintained under the same controlled environmental conditions without *P. triticina* inoculation. These conditions were used for disease severity assessment of Fielder WT, susceptible and sister-resistant controls, and prime edited *Lr67* lines.

## C) Supplemental Tables

**Table S1.**
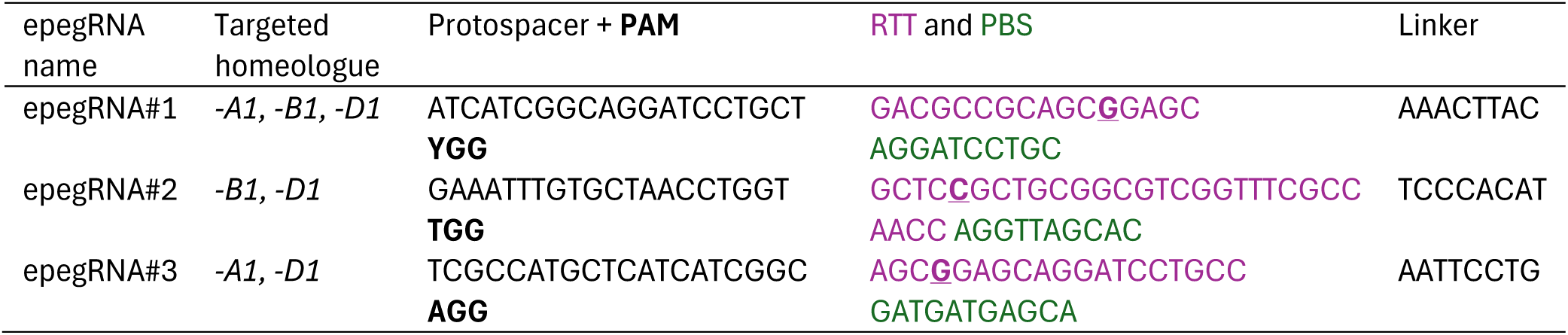
Sequences of epegRNAs used to install the TaSTP13G144R substitution. Includes protospacer, reverse transcription template (RTT), primer binding sequence (PBS), and linker. TaSTP13G144R G-to-C nucleotide is bold and underlined in the RTT.

**Table S2.** CAPS-based genotyping calls for *TaSTP13-A1*, *-B1*, and *-D1* across all T_0_ plantlets. “LostX” lines did not regenerate. Chim/Het: chimeric or heterozygous edit; Hom: homozygous edit.

| **Line ID** | ***TaSTP13-A1* CAPS** | ***TaSTP13-B1* CAPS** | ***TaSTP13-D1* CAPS** |
|---|---|---|---|
| Lost1 | Unedited | Unedited | Unedited |
| Lost2 | Chim/Het | Chim/Het | Chim/Het |
| Lost3 | Chim/Het | Chim/Het | Unedited |
| Lost4 | Chim/Het | Chim/Het | Unedited |
| Lost5 | Chim/Het | Chim/Het | Chim/Het |
| Lost6 | Chim/Het | Chim/Het | Unedited |
| T0-1 | Chim/Het | Chim/Het | Unedited |
| T0-2 | Chim/Het | Chim/Het | Unedited |
| T0-3 | Chim/Het | Chim/Het | Unedited |
| T0-4 | Chim/Het | Chim/Het | Chim/Het |
| T0-5 | Chim/Het | Chim/Het | Chim/Het |
| T0-6 | Chim/Het | Chim/Het | Chim/Het |
| T0-7 | Hom | Chim/Het | Chim/Het |
| T0-8 | Chim/Het | Chim/Het | Unedited |
| T0-9 | Chim/Het | Chim/Het | Unedited |
| T0-10 | Chim/Het | Chim/Het | Unedited |
| T0-11 | Chim/Het | Chim/Het | Chim/Het |
| T0-12 | Chim/Het | Hom | Chim/Het |
| T0-13 | Chim/Het | Chim/Het | Chim/Het |
| T0-14 | Hom | Hom | Hom |
| T0-15 | Chim/Het | Unedited | Unedited |
| T0-16 | Unedited | Unedited | Unedited |
| T0-17 | Chim/Het | Chim/Het | Chim/Het |
| T0-18 | Chim/Het | Chim/Het | Chim/Het |
| T0-19 | Chim/Het | Chim/Het | Chim/Het |
| T0-20 | Chim/Het | Chim/Het | Chim/Het |
| T0-21 | Chim/Het | Chim/Het | Chim/Het |
| T0-22 | Unedited | Unedited | Unedited |
| T0-23 | Hom | Hom | Chim/Het |
| T0-24 | Hom | Hom | Unedited |
| T0-25 | Chim/Het | Chim/Het | Unedited |

**Table S3.** Sanger sequencing genotype calls for *TaSTP13-A1*, *-B1*, and *-D1* in T_0_ plants. Genotypes were classified based on decomposition analysis using TIDER. Values in brackets indicate the estimated proportion of edited nucleotides at the target site and were used to assign genotypes: <5%: unedited; 5–40% chimeric edit; 40–80% heterozygous edit; >80% homozygous edit.

| **Line ID** | ***TaSTP13-A1* CAPS** | ***TaSTP13-B1* CAPS** | ***TaSTP13-D1* CAPS** |
|---|---|---|---|
| T0-1 | Heterozygous (70.1%) | Heterozygous (56.7%) | Chimeric (23.7%) |
| T0-2 | Heterozygous (59.4%) | Heterozygous (54.9%) | Chimeric (21.3%) |
| T0-3 | Heterozygous (43.6%) | Chimeric (39.5%) | Unedited (4.2%) |
| T0-4 | Homozygous (98.3%) | Heterozygous (76.7%) | Heterozygous (40.2%) |
| T0-5 | Homozygous (88.6%) | Heterozygous (75.7%) | Chimeric (39.8%) |
| T0-6 | Homozygous (98.3%) | Heterozygous (69%) | Chimeric (28.6%) |
| T0-7 | Homozygous (98.3%) | Homozygous (98.2%) | Heterozygous (47.5%) |
| T0-8 | Heterozygous (54%) | Heterozygous (57.2%) | Chimeric (13.8%) |
| T0-9 | Heterozygous (63%) | Heterozygous (43.6%) | Chimeric (14%) |
| T0-10 | Heterozygous (59.6%) | Heterozygous (48.2%) | Chimeric (21.6%) |
| T0-11 | Heterozygous (74.4%) | Homozygous (96.7%) | Chimeric (26.6%) |
| T0-12 | Homozygous (97.9%) | Homozygous (99.5%) | Chimeric (39.3%) |
| T0-13 | Homozygous (97.8%) | Homozygous (99.5%) | Heterozygous (50.7%) |
| T0-14 | Homozygous (95.6%) | Homozygous (99.5%) | Homozygous (98.9%) |
| T0-15 | Chimeric (35%) | Chimeric (24%) | Unedited (1.3%) |
| T0-16 | Unedited (0%) | Unedited (0.1%) | Unedited (0%) |
| T0-17 | Homozygous (98.4%) | Heterozygous (78.1%) | Heterozygous (57.4%) |
| T0-18 | Homozygous (84.9%) | Homozygous (99.5%) | Heterozygous (74%) |
| T0-19 | Heterozygous (77.9%) | Homozygous (98.8%) | Heterozygous (71.9%) |
| T0-20 | Heterozygous (75.9%) | Homozygous (99.5%) | Heterozygous (66.7%) |
| T0-21 | Homozygous (81.5%) | Heterozygous (69.7%) | Chimeric (28.7%) |
| T0-22 | Unedited (0%) | Unedited (0%) | Unedited (0.2%) |
| T0-23 | Homozygous (98.4%) | Homozygous (99.5%) | Heterozygous (72.9%) |
| T0-24 | Homozygous (98.4%) | Homozygous (99.5%) | Chimeric (29.8%) |
| T0-25 | Heterozygous (62.3%) | Heterozygous (40.7%) | Chimeric (12.2%) |

**Table S4.** Zygosity (%) calls for TaSTP13-A1, -B1, and -D1 in T3 plants.

|  | Sample | A-Genome | B-Genome | D-Genome | Treatment |
|---|---|---|---|---|---|
| **A-Edit** | 1 | 95 | 6 | 13 | Infected |
|  | 2 | 95 | 1 | 7 | Mock |
|  | 3 | 88 | 7 | 0 | Infected |
|  | 4 | 85 | 0 | 9 | Mock |
|  | 5 | 96 | 8 | 0 | Infected |
|  | 6 | 96 | 6 | 0 | Mock |
|  | 7 | 96 | 0 | 0 | Infected |
|  | 8 | 96 | 0 | 0 | Mock |
| **AB-Edit** | 1 | 96 | 94 | 9 | Infected |
|  | 2 | 95 | 73 | 0 | Mock |
|  | 3 | 96 | 86 | 0 | Infected |
|  | 4 | 85 | 94 | 0 | Mock |
|  | 5 | 95 | 93 | 0 | Infected |
|  | 6 | 95 | 93 | 0 | Mock |
|  | 7 | 95 | 87 | 0 | Infected |
|  | 8 | 93 | 92 | 0 | Mock |
|  | 9 | 95 | 92 | 5 | Infected |
| **ABD-Edit** | 1 | 95 | 91 | 77 | Infected |
|  | 2 | 96 | 92 | 74 | Mock |
|  | 3 | 95 | 93 | 70 | Infected |
|  | 4 | 95 | 92 | 71 | Mock |
|  | 5 | 95 | 93 | 80 | Infected |
|  | 6 | 95 | 93 | 70 | Mock |
|  | 7 | 96 | 93 | 70 | Infected |
|  | 8 | 96 | 94 | 70 | Mock |
|  | 9 | 95 | 92 | 71 | Infected |
|  | 10 | NA | 92 | 75 | Mock |

**Supplemental Table 5.**
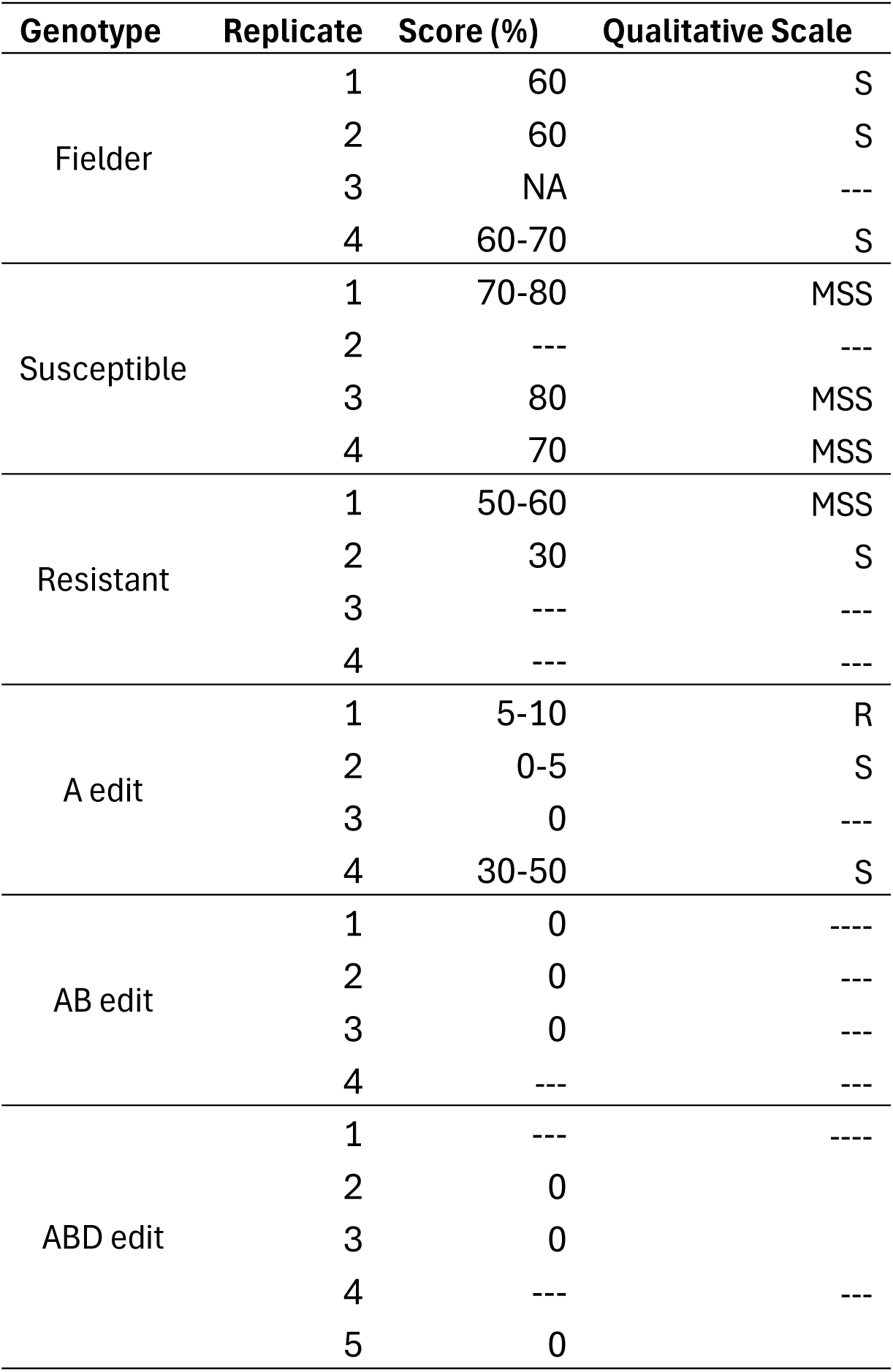
Semi quantitative rusts scoring at 13 days post inoculation. For qualitative scale, S = Susceptible, MSS = Moderate susceptible, R = Resistance.

**Supplemental Table 6.**
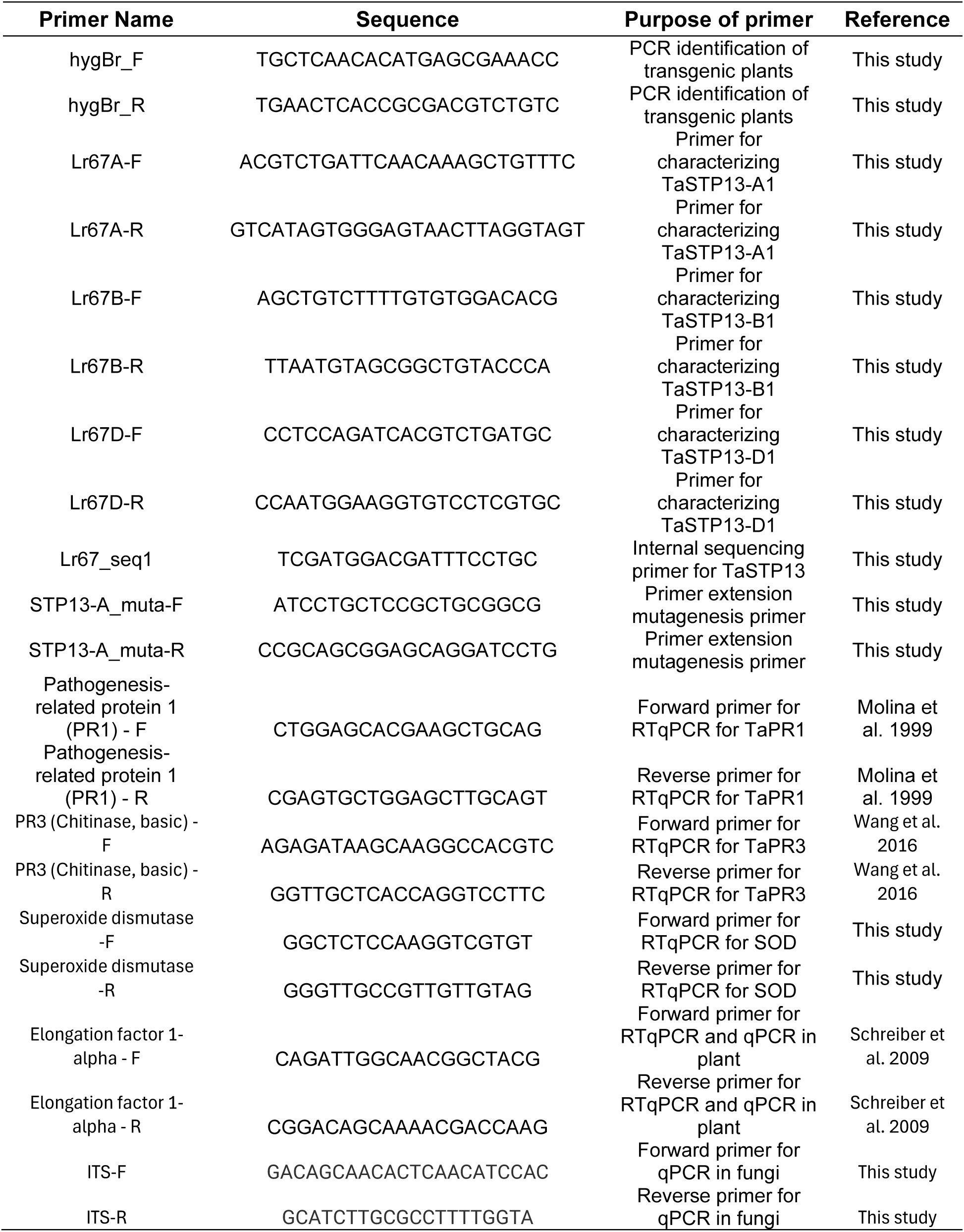
Primers used in this study for the characterization of plants and primer to generate gene edited lines in *TaSTP13*.

### D) Supplemental sequences

**Supplemental Sequence 1.** Synthesized fragment of the three epegRNA expression cassettes that install the G144R substitution in all homoeologs of *TaSTP13*.

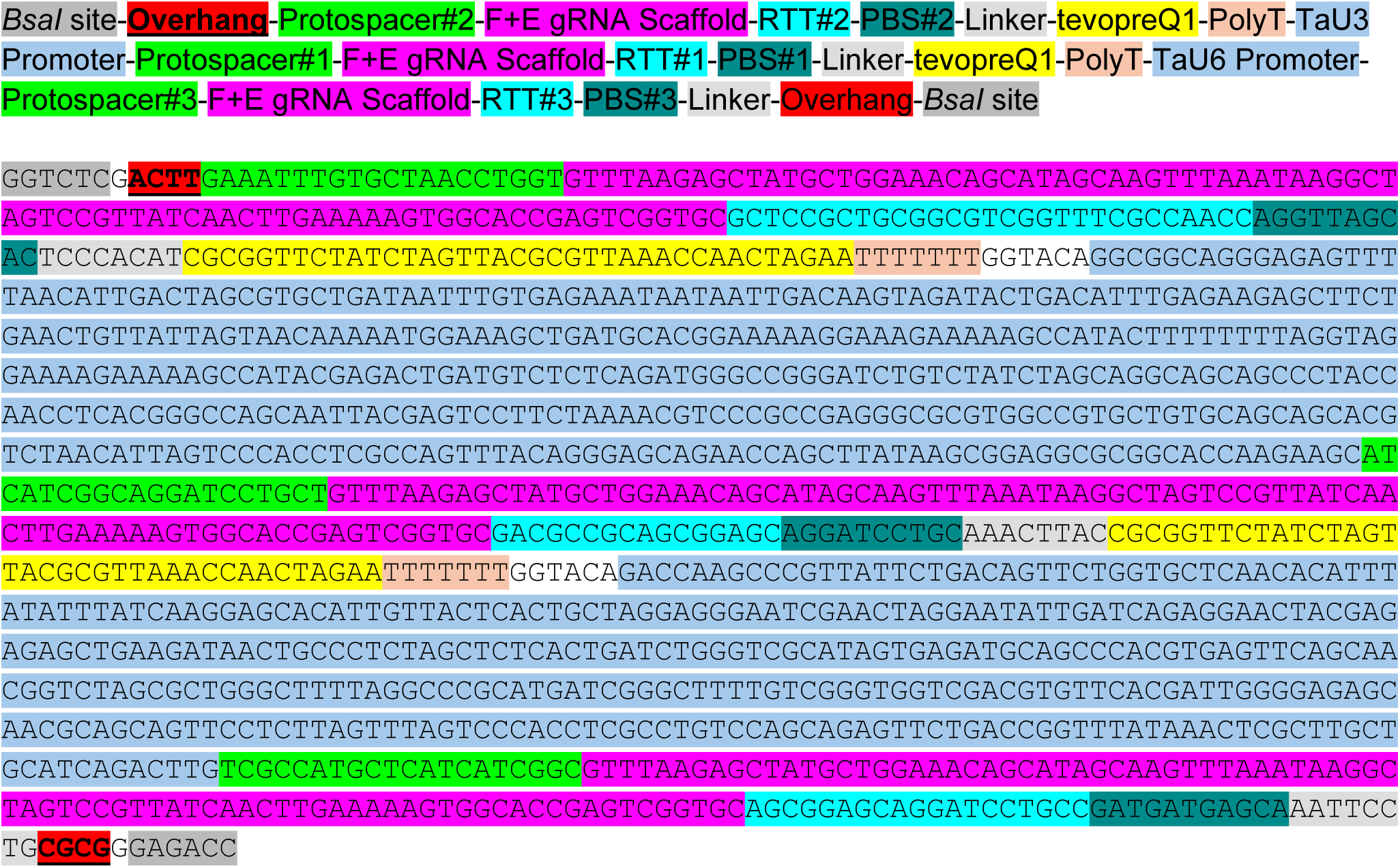

**Supplemental Sequence 2.** Synthesized fragment for the construction of JWPE1.

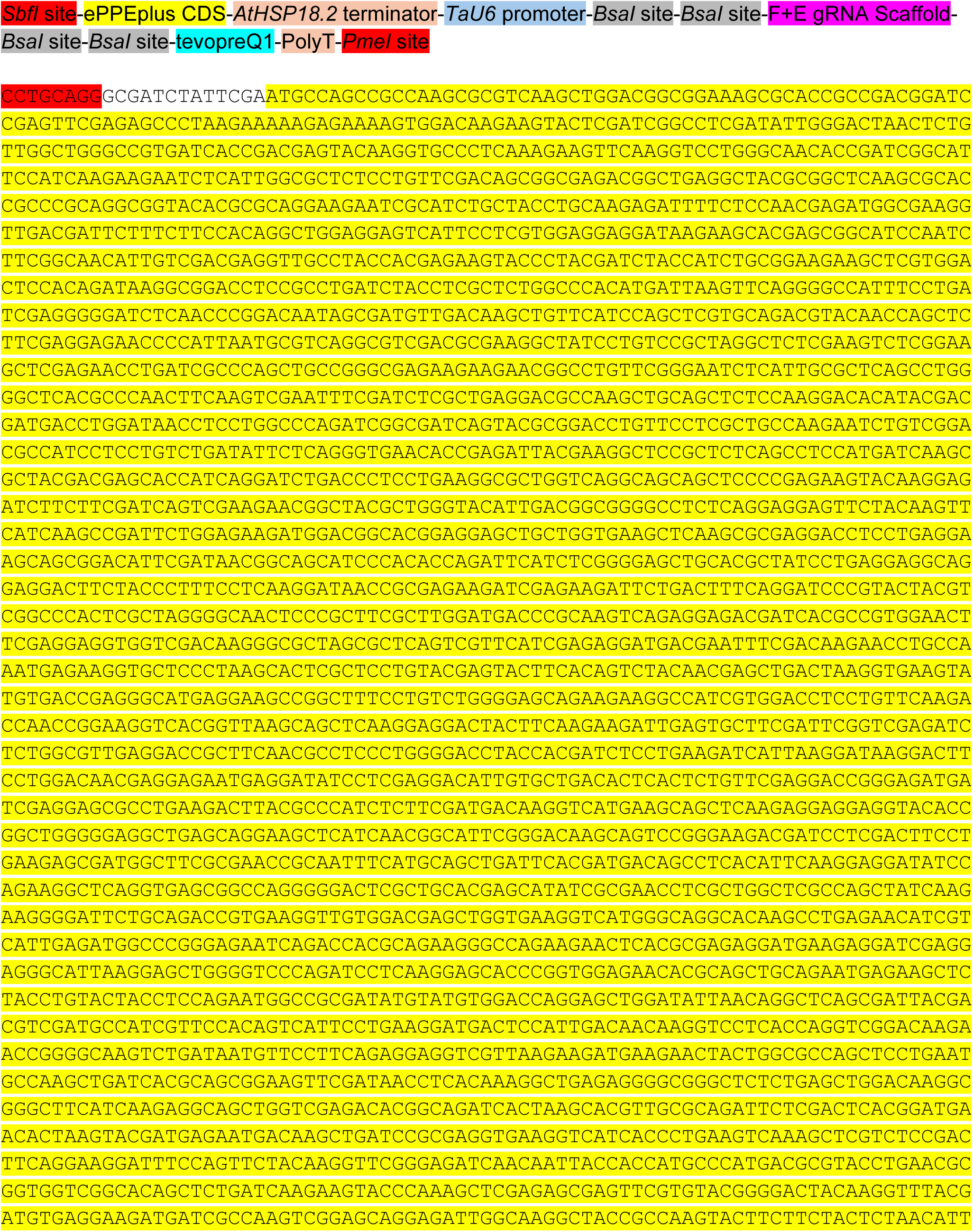
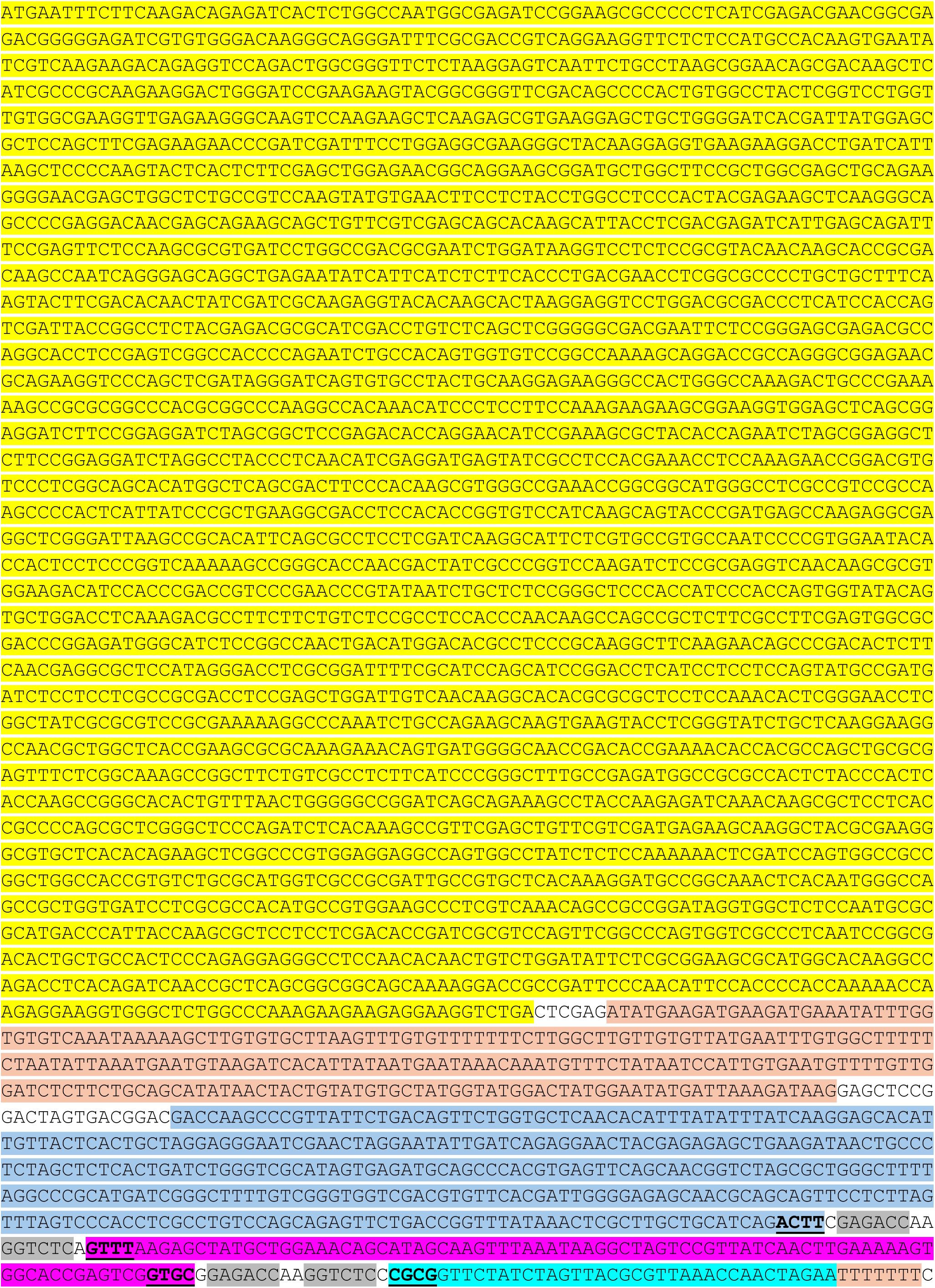
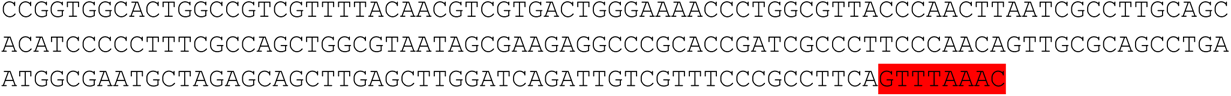

**Supplemental Sequence 3.** ePPEplus expression cassette within JWPE1.

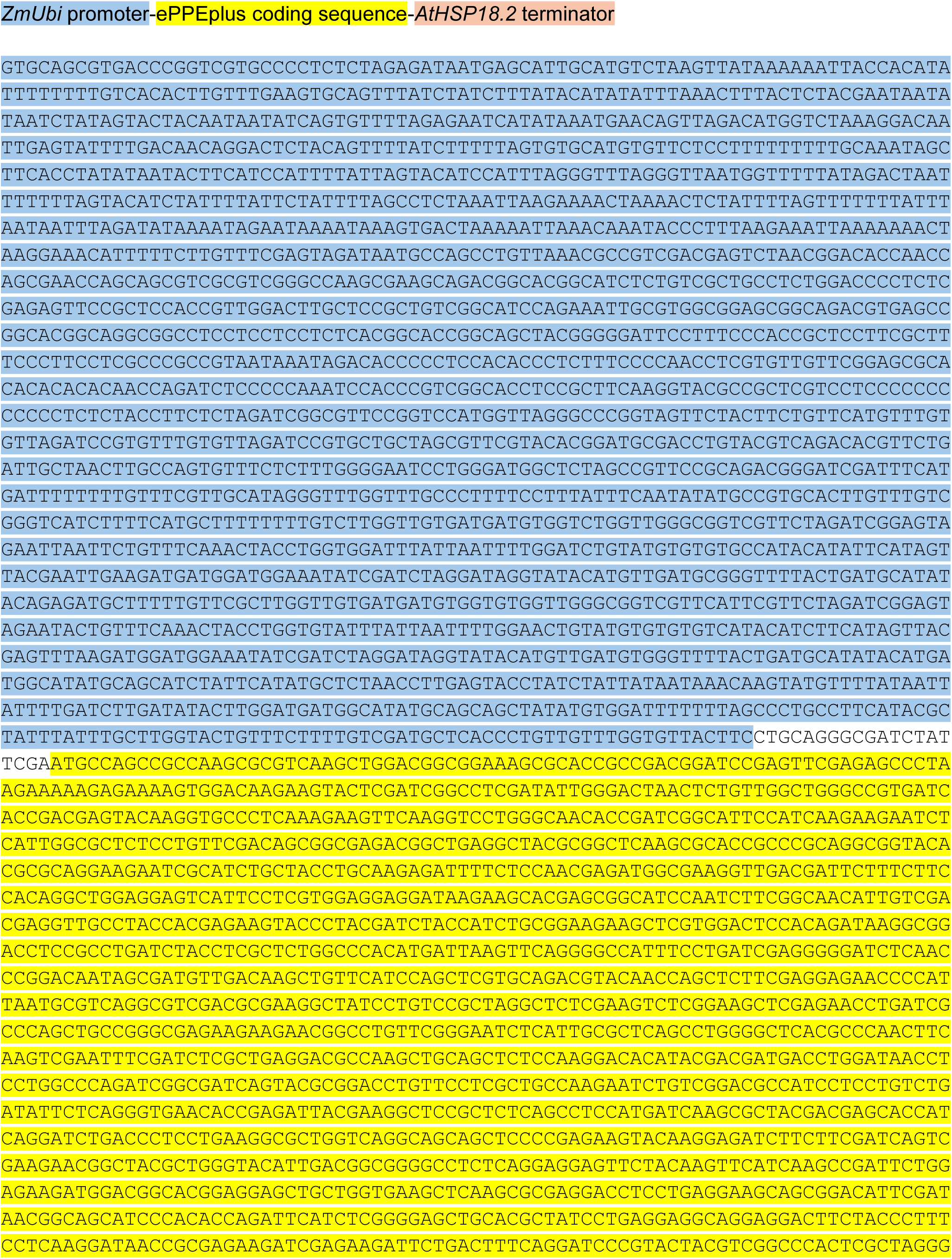
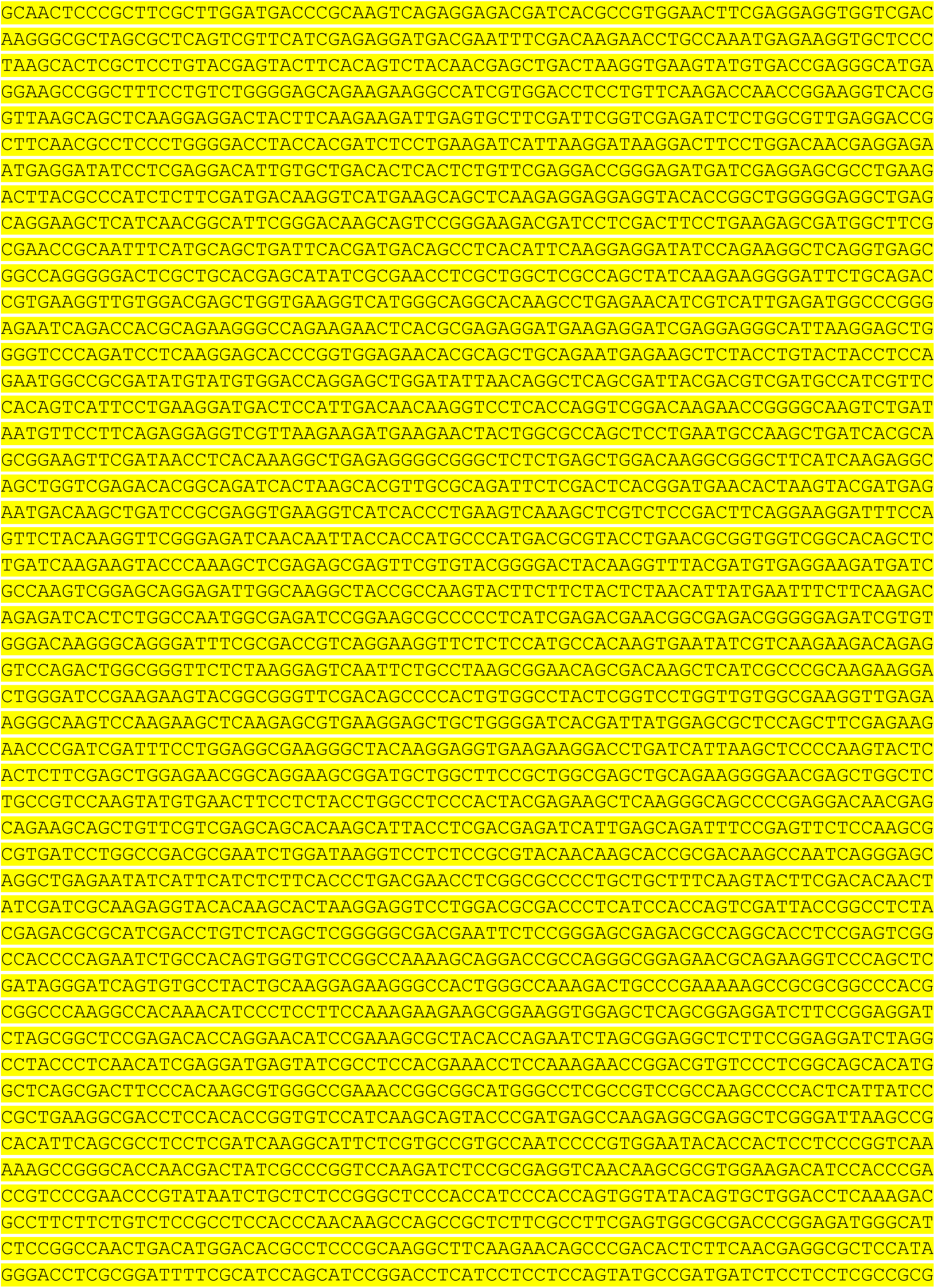
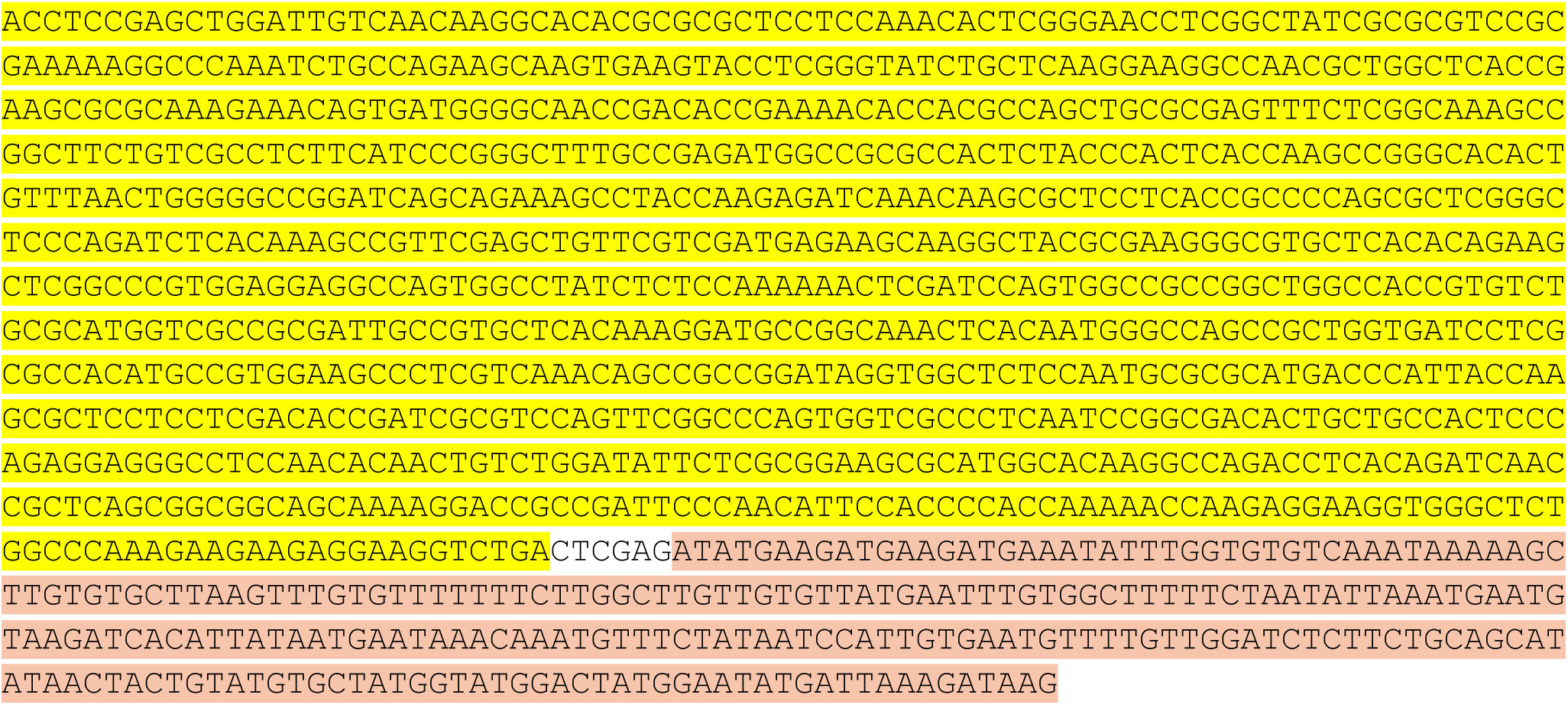

**Supplemental Sequence 4.** epegRNA expression cassette within JWPE1.

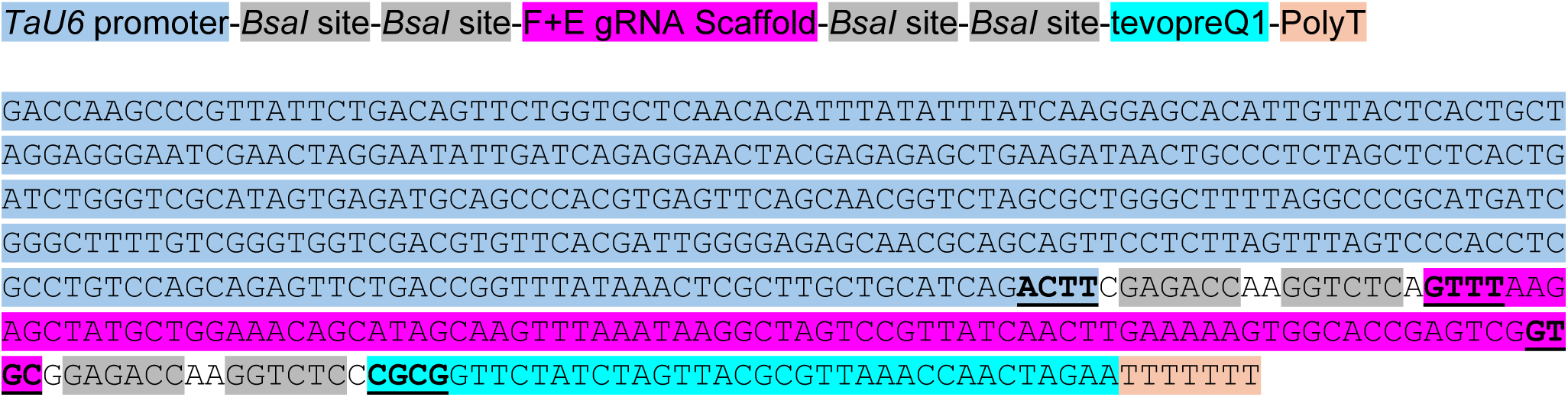

**Supplemental Sequence 5.** Chromatograms of all T3 prime-edited wheat plants used for *Puccinia triticina* inoculation and mock treatments

All sequence data alignment is available at DOI: 10.6084/m9.figshare.33959875

## Notes

https://doi.org/10.6084/m9.figshare.33959875

### Competing Interest Statement

The authors have declared no competing interest.

